# ATG7 function promotes pancreatic cancer progression independently of autophagy

**DOI:** 10.64898/2026.06.01.729197

**Authors:** Clémence Larat, Adrian Lopez Garcia de Lomana, Valgerdur Hjaltalin, Margret H Ogmundsdottir

## Abstract

Autophagy is a cellular degradation process that recycles dysfunctional components to maintain cellular homeostasis. Beyond this canonical role, autophagy-related proteins, such as the essential autophagy initiation protein ATG7, are increasingly recognized to have autophagy independent functions in diverse biological processes and disease contexts, including cancer progression and metastasis. However, the mechanisms underlying these autophagy independent functions remain unclear. Previously, we identified a short isoform, ATG7(2), that lacks canonical autophagy activity. To understand the unique role of ATG7(2), we analysed clinical data from publicly available databases and found that high *ATG7(2)* expression is associated with poor prognosis in pancreatic adenocarcinoma (PAAD). Using CRISPR/Cas9 in PAAD cells, we selectively knocked out the canonical isoform ATG7(1) or total ATG7. While total knock-out of ATG7 slowed proliferation and migration of PAAD cells, high levels of ATG7(2) were found to enhance both processes. In addition, RNA sequencing linked ATG7(2) with immune signalling, extracellular matrix organization and cell-cell interactions. Critically, ATG7(2) inhibition in a murine xenograft model substantially reduces tumour growth and overall progression *in vivo*, establishing functional relevance in a physiological tumour context. Together, these results suggest that ATG7(2) has a role in regulating immune signalling in PAAD cells, contributes to migration and proliferation in an autophagy-independent manner, and suggests ATG7(2) as a potential therapeutic target for the treatment of PAAD.

## INTRODUCTION

Macro autophagy, hereafter referred to as autophagy, is a cellular degradation process that recycles damaged organelles and other intracellular components to maintain cellular homeostasis [1], [2]. Autophagy is initiated primarily at the endoplasmic reticulum, where a phagophore forms around cargo and subsequently fuses with lysosomes for degradation [2]. In cancer, autophagy plays context-dependent roles, acting as both a tumour suppressor by limiting damage accumulation and a tumour protector by supporting proliferation and drug resistance in established tumours [3].

Essential to this initiation step is the key autophagy protein ATG7, which drives phagophore elongation through the lipidation of ATG8 family proteins [4]. Both autophagy and ATG7 specifically have been identified as prognosis markers in diverse cancer types [3], [5], [6]. ATG7 has also been shown to regulate tumour immunity by influencing the infiltration of immune cells within the tumour microenvironment, including CD8+ cells, which are part of the anti-tumoral immune response [7], [8]. However, the role of ATG7 in tumoral immunity is not fully understood and appears to be dependent on tumour type, as ATG7 inhibition has also been shown to promote anti-tumoral immune responses in certain cancers, including colorectal cancer [9], [10]. Immune escape and tumour immunity in general have been identified as key modulators of tumour progression and drug resistance [11], [12], [13]. ATG7 has also been linked with pancreatic adenocarcinoma (PAAD), in which high ATG7 expression is associated with poor prognosis and rapid cancer progression [14], [15]. Additionally, ATG7 has been shown to play an autophagy-independent role in pancreatic ductal adenocarcinoma (PDAC), the most prevalent subtype of PAAD [16]. Together, these observations suggest that ATG7 may influence tumour progression through mechanisms beyond its canonical role in autophagy.

A short isoform of ATG7, termed ATG7(2), was previously identified and lacks the ability to carry out the canonical role of ATG7 in autophagy initiation [17]. While ATG7(1) (703 amino acids, UniProt O95352-1) contains all exons and is responsible for the canonical activity of the protein, ATG7(2) (676 amino acids, UniProt O95352-2) lacks an exon, number 17, resulting in a 27 amino acid truncation in the C-terminal region and preventing lipidation of ATG8 proteins [17]. Although ATG7(2) has been shown to bind metabolic proteins [18], the functional role of ATG7(2), or its role in disease progression, remains unclear.

To explore the potential role of ATG7(2) in PAAD progression and metastasis, we used CRISPR/Cas9 technology in human PAAD Capan-1 cells to selectively knock out ATG7(1) or total ATG7. *In vitro* phenotyping and characterization of these cells revealed that ATG7(2) drives PAAD cell proliferation and migration and regulates the expression of genes involved in immune pathways, extracellular matrix (ECM) organization, and cell–cell interactions. Furthermore, ATG7(2) was found to be involved in cytokine secretion, and its inhibition *in vitro* led to decreased cell proliferation and migration, as well as reduced expression and secretion of various cytokines. Finally, formation of xenografts from CRISPR-engineered PAAD cells in BALB/c nude mice, combined with intratumoral siRNA treatment, revealed that ATG7(2) inhibition efficiently reduced tumour progression *in vivo*. These findings establish ATG7(2) as a regulator of tumour progression independent of autophagy and position it as a potential therapeutic target in PAAD.

## MATERIAL AND METHODS

### Cell culture

Capan-1 cells were obtained from the Pathology department at Landspitali, University of Iceland. They were cultivated on collagen rat tail I (Sigma C3867) coated vessels in IMDM (Gibco 12440053) supplemented with 20% FBS (Gibco A5256701) in an incubator at 37°C with 5% CO_2_. Panc 08.13 cells were obtained from the Cancer Research UK Scotland Institute. They were cultivated in RPMI (Gibco 11875093) supplemented with 15% FBS (Gibco A5256701) and 10 units/mL of human recombinant insulin (SAFC 91077C). DMEM without Glucose (Gibco A1443001) was used to induce starvation while DMEM high Glucose with 20% FBS (Gibco 11995073) was used on the control condition.

### CRISPR/Cas9

#### gRNA and Cas9 transfection

For each reaction, 1μL of 100uM crRNA and 1μL of 100uM tracrRNA (IDT Alt-R CRISPR-Cas9 5’ ATTO 550 #1077024) were mix in a final volume of 100μL of nuclease free duplex buffer (IDT #1072570) and incubated at 95°C for 5 minutes to generate gRNAs. 1,6μL of Cas9 (IDT Alt-R S. p. Cas9 #1081059) was diluted in a final volume of 100μL in OptiMEM (Gibco 11058021). In the well of a 24-well plate coated with collagen, 88μL OptiMEM, 6μL gRNA and 6μL diluted Cas9 were mixed and incubated at room temperature for 5 minutes to generate the RNP complex. 99μL of OptiMEM and 1μL of RNAiMax (Invitrogen 13778) were added to the RNP complex and incubated 20 minutes at room temperature. 1×10^4^ – 2×10^4^ cells were added to the well in a final volume of 500μL IMDM 20% FBS and grown at 37°C with 5% CO_2_. Control reaction was performed with DEPC water instead of crRNA.

The following sequences were targeted by the crRNA:

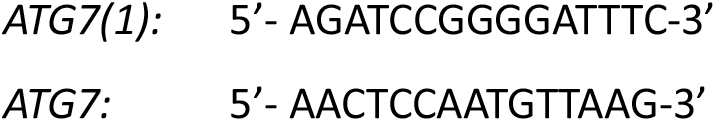

#### Single cell cloning

After 48 hours, the cells were suspended and diluted to a concentration of 1 – 2 cells/mL. Then, 7mL of cell suspension was added to 10cm tissue dishes coated with collagen, and 24 dishes were used per CRISPR reaction. The cells were grown at 37°C with 5% CO_2_ for 17 days and 5 tracrRNA positive clones were picked from each dish and transferred to a collagen-coated well on a 24 well plate. The 120 clones from each reaction were grown until confluent.

#### DNA extraction

Cell pellets from confluent clones were collected and resuspended in 30μL of Igepal buffer (52,5mM Tris-HCl; 52,5mM KCl; 3,3075mM MgCl_2_; 0,26125% Igepal; 0,525% Tween 20; 0,025% proteinase K (NEB P8107)) and heated for 90 minutes at 60°C, followed by 15 minutes at 95°C. Samples were centrifuged for 15 minutes at 16 000g and 20μL of the supernatant were collected. DNA was quantified using NanoDrop One Spectrophotometer (ThermoFisher) and samples were diluted to a final concentration of 50ng.μL^−1^ in DEPC water.

#### PCR reaction

For each reaction, 0,5μL of 10µM forward primer, 0,5μL of 10µM reverse primer, 1,5μL DEPC water, 12,5μL OneTaq 2X Master Mix with standard buffer (NEB #M0482) and 10μL of 50ng.μL^−1^ were mixed and underwent the following program: 94°C 30sec – (94°C 30sec – 45°C 60sec – 68°C 60sec) x30 – 68°C 5 minutes.

The following primers were used:

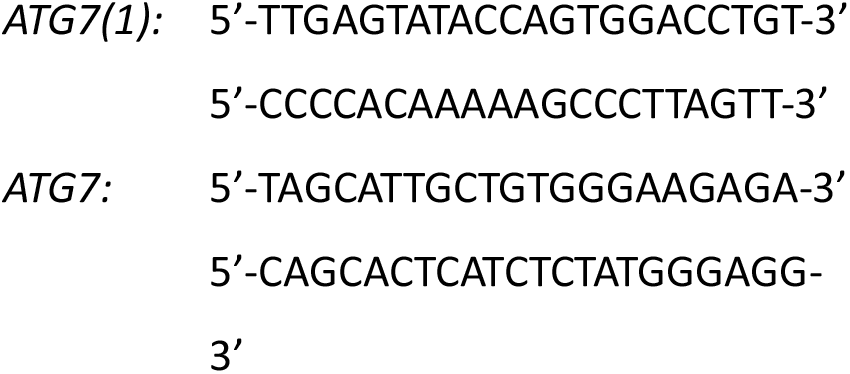

#### Sanger sequencing

The PCR products were cleaned-up using ExoSAP-IT (ThermoFisher #78200.200.ML): 0,5μL ExoSAP was added to each 5μL of PCR product and heated at 37°C for 30 minutes and 90°C for 10 minutes. For each sequencing reaction, 3,2μL of BigDye Direct Sequencing Master Mix (Applied Biosystems 4458687), 0.8μL DEPC water, 0,5μL sequencing primer and 0,5μL ExoSAP product were mixed and underwent the following program: 95°C 3min – (95°C 20sec – 55°C 7sec – 60°C 4min) x30. Sequencing was read by DeCode Genetics, Inc. and results were compared with WT ATG7 sequence using Benchling Biology Software.

The following sequencing primers were used:

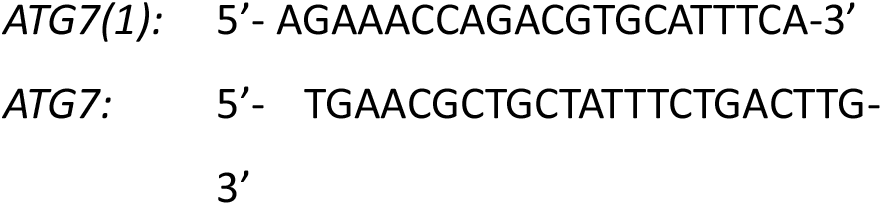

### Quantitative Real-Time PCR

Total RNA was extracted from cells using the Quick-RNA Miniprep Kit (Zymo Research R1055) following the manufacturer’s protocol. For tumour samples, the tissue was resuspended in DNA/RNA shield buffer (Zymo Research R1100) using a micropestle before isolating total RNA with the Quick-RNA Miniprep Kit. RNA samples were diluted to a final concentration of 100ng.μL^−1^ in nuclease-free water. Reverse transcription was performed using the High-Capacity cDNA Reverse Transcription Kit (Applied Biosystems no. 4368814) following the manufacturer’s protocol for 10μL reaction volume. cDNA samples were diluted 1:50 in nuclease-free water. qPCR was performed using Luna Universal qPCR Master Mix (New England Biolabs Inc., #M3003) following the manufacturer’s protocol for 10μL reaction volume with 4μL of cDNA. The data weas normalized to actin mRNA. The following primers were used at a final concentration of 0,25μM:

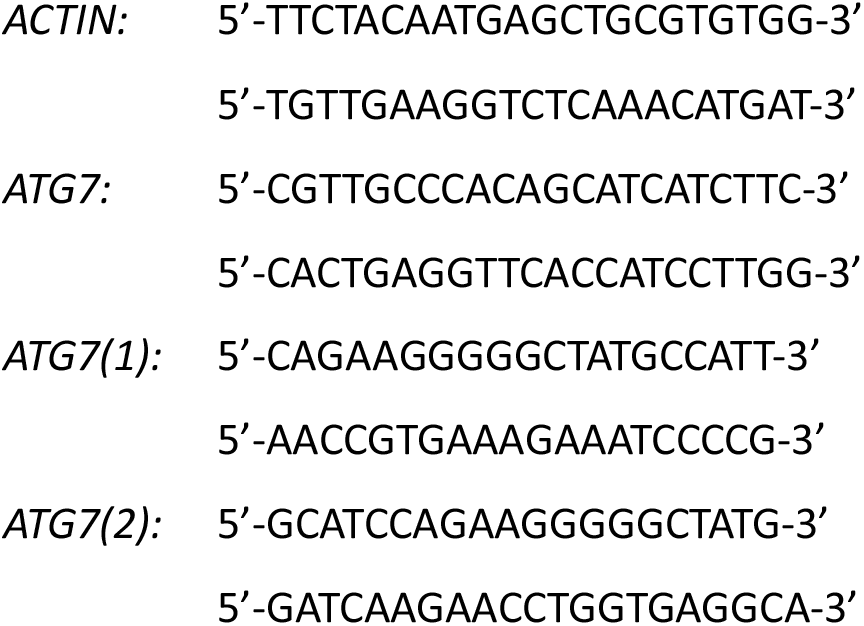

### Western Blotting

Samples were collected in 1X RIPA buffer (Abcam ab156034) with 1% protease inhibitor (ThermoFisher 1861281) or through another specified method. Protein loading dye (NEB B7703S) was added to the samples following manufacturer’s instructions and the samples were boiled for 5 min at 95°C. Proteins were separated by SDS–polyacrylamide gel electrophoresis (SDS-PAGE) at 150V on 12% acrylamide gels. Then, the proteins were transferred onto PVDF membranes (Thermo Fisher Scientific, 88520) at 25V overnight or 400mA for 2h on ice. Membranes were blocked in 5% BSA (Sigma, A9647) in TBS with 0,1% Tween-20 (Sigma, P1379) for 30min at RT. Primary antibodies were incubated overnight at 4°C or 3h at room temperature. Finally, DyLight 800 anti-mouse and DyLight 680 anti-rabbit (Cell Signaling Technology) were applied for 45min at RT. All the anti-bodies were used at the dilutions recommended by manufacturers. Odyssey imaging system and Image Studio version 2.0 (LI-COR Biosciences) were used to scan the blots.

**Table 1:** Primary antibodies used for Western-Blotting.

| Target | Species | Dilution | Manufacturer |
| --- | --- | --- | --- |
| Actin | Mouse | 1:10000 | Millipore, MAB1501 |
| GAPDH | Rabbit | 1:5000 | Cell Signaling Technology, #5174 |
| H3 | Rabbit | 1:2000 | Cell Signaling Technology, #4499 |
| ATG7 | Rabbit | 1:1000 | Cell Signaling Technology, #8558 |
| LC3B | Rabbit | 1:1000 | Cell Signaling Technology, #2775 |
| p62 | Mouse | 1:1000 | Cell Signaling Technology, #5114 |
| Ajuba | Rabbit | 1:1000 | Cell Signaling Technology, #34648 |
| YAP | Mouse | 1:1000 | Cell Signaling Technology, #12395 |
| SHP2 | Mouse | 1:2000 | Invitrogen, MA5-17160 |

### Cell fractionation

Cells were collected with 1X trypsin and centrifuged for 5min at 600g. Supernatant was discarded, and the pellet was washed with 200μL of 1X ice-cold PBS and centrifuged for 5min at 600g. On ice, cells were resuspended in 200μL of swelling buffer (10mM HEPES pH7,9; 1,5mM MgCl_2_; 10mM KCl; 0,5mM DTT; 1% protease inhibitor) and centrifuged for 5min at 600g and 4°C. The supernatant was discarded, and the pellet was resuspended in 200μL of swelling buffer, spined 5min at 600g and 4°C and the supernatant was discarded. Cells were resuspended in 100μL of lysis buffer (10mM HEPES pH7,9; 1,5mM MgCl_2_; 10mM KCl; 0,5mM DTT; 0,1% NP-40; 1% protease inhibitor) and incubated on ice for 10min. 30μL of cell lysate were collected and 3μL of 10% Triton were added. The remaining lysate was centrifuged for 5min at 3000g and 4°C and the supernatant containing the cytoplasmic fraction was collected. The pellet was washed with 150μL of 1X ice-cold PBS and centrifuged for 5min at 3000g and 4°C. The supernatant was discarded and the pellet was resuspended in 70μL of lysis buffer supplemented with 0,25U.mL^−1^ benzonase nuclease and 1% Triton. Protein concentration was determined using BCA assay (ThermoFisher 23227) and the samples were subsequently used for western blotting.

### Proliferation and migration assays

#### Proliferation assay

Capan-1 cells were seeded at a density of 2000 cells per well on a 96 well plate coated with collagen Rat tail I and placed in the IncuCyte Live-Cell imaging system. For knock-down, cells were reverse transfected with siRNA when seeded (see siRNA knock-down) and treatment was repeated with forward transfection every 72 hours. A single picture was taken of each seeded well with a 4x zoom every 4 hours over 11 days. For each well, cell confluency was normalized to the confluency of the first image (0h) directly in the IncuCyte 2022A Rev1 software. For each condition, 5 technical replicates were performed and the experiment was repeated 3 times for statistical analysis.

#### Migration assay

Capan-1 cells were seeded at a density of 35000 cells per well on a 96 well plate (Sartorius IncuCyte ImageLock 4379) coated with collagen Rat tail I and incubated for 24h at 37°C with 5%CO_2_. For knock-down, cells were reverse transfected with siRNA when seeded (see siRNA knock-down). Wounds were created using a woundmaker (Sartorius 4563). The pins of the woundmaker were washed 5min in sterile water and 5min in ethanol 70%. The plate was placed on the woundmaker and the cover was removed before placing the pins in the wells. The switch of the woundmaker was pressed down and the pins were removed from the wells without releasing the switch. The cells were rinsed once with PBS and treatment was added to the wells if applicable. The pins were subsequently washed 5min in Alconox 0,5%; 5min in Virkon S 1%; 5min in sterile water and 5min in ethanol 70%. The plate was placed in the IncuCyte Live-Cell imaging system. Two pictures were taken of each seeded well with a 10x zoom every 2 hours over 3 days. For each well, relative wound density was calculated directly in the IncuCyte® 2022A Rev1 software. For each condition, 5 technical replicates were done and the experiment was performed 3 times for statistical analysis.

#### siRNA knock-down

Cells were reverse- or forward-transfected with siRNA using OptiMEM (Gibco 11058021) and Lipofectamine RNAiMax transfection reagent (Invitrogen 13778) following manufacturer’s recommendation. The following siRNAs were used:

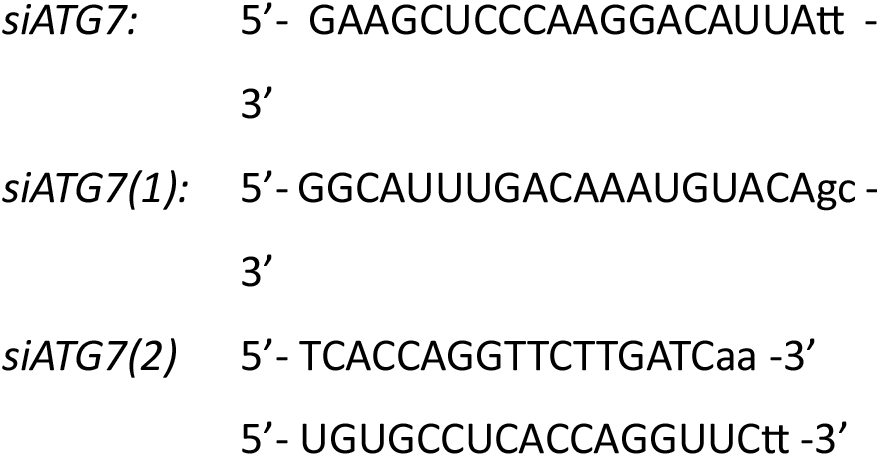

#### RNA-sequencing

Total RNA was extracted from cells using the Quick-RNA Miniprep Kit (Zymo Research R1055) following the manufacturer’s protocol and subsequently sequenced. We cleaned original FASTQ files using fastp version 1.0.1 using default parameters [19]. Next, we quantified gene expression from generated FASTQ files using kallisto version 0.51.1 [20] and the `--rf-stranded` option against the Ensembl Homo sapiens reference transcriptome (version 108). We generated a final table of transcriptome quantification at gene level and TPM units using the DESeq2 [21] function `tximport`. Then, we used DESeq2 version 1.26.0 [21] to determine statistically significant differentially expressed genes across experimental conditions (Benjamini– Hochberg correction α = 0.05; adjusted *P* < 0.05). Prior to statistical testing, very lowly expressed genes were excluded (less than 10 counts across at least three samples). We considered response genes that displayed expression differences of more than 50 normalized counts, relative expression differences larger than one log_2_ fold change and statistical significance (adjusted *P* < 0.05). The GitHub repository https://github.com/ClemLarat/ATG7_PAAD contains the computational tools to reproduce the quantitative sequence analysis results presented in this work.

#### Immunofluorescence

1×10^4^ cells were seeded in the well of a 12-well chamber slide (Ibidi #81201) coated with collagen. After 24h, culture medium was removed, the cells were covered in 2-3mm of formaldehyde 4% and incubated 15min at room temperature. The samples were washed 3 times for 5min in 1X PBS before being blocked at room temperature for 1h in 100µL of 1X PBS, normal goat serum 5%, Triton X-100 0,3%. Blocking solution was removed and samples were incubated overnight at 4°C in 100µL of primary antibody solution (1X PBS, BSA 1%, Triton X-100 0,3%, mAb 1:100). The samples were washed 3 times 5min in 1X PBS before incubating 2h at room temperature protected from light in 100µL of secondary antibody solution (1X PBS, BSA 1%, Triton X-100 0,3%, mAb 1:500). The samples were washed 3 times for 5min in 1X PBS before removing the chambers from the slide and mounting the coverslip with DAPI fluoromount-G (Invitrogen #00-4959-52). The samples were subsequently stored at 4°C protected from light before imaging on an Olympus FV4000 confocal microscope.

**Table 2:** Antibodies used for immunostaining.

| Type | Target | Species | Dilution | Manufacturer |
| --- | --- | --- | --- | --- |
| Primary | ATG7 | Rabbit | 1:100 | Abcam, ab52472 |
|  | YAP | Rabbit | 1:50 | Cell Signaling Technology, #14074 |
| Phalloidin | F-Actin | n.a. | 1:1,000 | Abcam, ab176759 |
| Secondary (Alexa Fluor) | Mouse | Goat | 1:500 | Thermo, A21235 |
|  | Rabbit | Goat | 1:500 | Thermo, A11008 |

#### Animal experiments

All animals were housed in a pathogen-free environment with a 12-h light/dark cycle with access to food and water ad libitum. Female BALBc/nude mice were acquired from Janvier labs at 4 weeks of age. Experiments were designed and carried out in compliance with the Icelandic Food and Veterinary Authority (Matvælastofnun, project license 24071280). Mice were randomized in 6 cohorts of 5 animals each. At 6 weeks of age, Capan-1 cells were counted and resuspended in PBS at a concentration of 1.5×10^7^cells/mL, Matrigel (Corning 356234) was added to the cell suspension at a 1:1 ratio and 100µL were injected subcutaneously in the right flank of each mouse using a 1.5mL pipette with a 27gauge needle. Mice were subjected to weekly weighing and when detectable, the tumours were measured weekly using a calliper, and the volume was calculated as follows: 0.5236 x Length (mm) x Width (mm) x Height (mm) = Volume (mm^3^). On the third week after xenograft, the animals were treated by intra-tumoral injections of siRNA (Ambion *In Vivo* siRNA, ThermoFisher Scientific) in lipid nanoparticles (*in vivo*-jet PEI, Polyplus Sartorius), as per manufacturers recommendation for intra-tumoral delivery. The treatment was repeated twice weekly over the course of 5 weeks. 72h after the last intratumoral injection, the animals were euthanized by inhalation of anaesthesia gas (vetfluorane), and secondary cervical dislocation. Animals were weighed and their tumours were measured. Tumours were subsequently extracted, weighed and kept on ice before all tumours were photographed together. Tumour samples were placed in DNA/RNA shield (Zymo Research R1100) and stored at 4°C before subsequent RNA isolation.

#### Statistical analysis

All experiments were performed in at least three biological replicates. All statistical tests were performed in GraphPad (Prism), except for RNA-seq analysis, which was performed in R-studio. Student t-test was used for comparison between 2 groups only, one-way and two-way ANOVA with Fisher’s LSD test were used to compare 3 groups or more, and Pearson correlation coefficient was used for correlation analysis.

## RESULTS

### ATG7(2) is linked with poor prognosis in PAAD, and is involved in immunity

Alternative splicing of *ATG7* pre-mRNA leads to the expression of two main isoforms termed *ATG7(1)* and *ATG7(2)* (Fig. 1.A). To analyse the association of *ATG7* isoforms with cancer, we downloaded survival and RNA expression data of cancer patients from TCGA database and analysed it on the Gepia webpage [22]. Results showed that high *ATG7* expression is significantly associated with poor prognosis in Kidney chromophobe (KICH, p = 0.043), Liver hepatocellular carcinoma (LIHC, p = 0.014) and Pancreatic adenocarcinoma (PAAD, p = 0.036), while it is significantly associated with good prognosis in Kidney renal clear cell carcinoma (KIRC, p = 0.007) (Fig. 1.B). Further analysing this trend for *ATG7(1)* and *ATG7(2)* isoforms specifically, we observed a negative association of high *ATG7(1)* expression with survival in LIHC, a positive association of high *ATG7(2)* expression with survival in KIRC and a negative association in PAAD (Fig. 1.B). Kaplan-Meier plots of this data further showed that the association of high *ATG7* expression with poor prognosis in PAAD is likely through *ATG7(2)* (Fig. 1.C). RNA expression data from GTEx was downloaded and expression of *ATG7* isoforms was compared between pancreatic tissue and PAAD samples (Fig. 1.D) and showed increased expression of both isoforms in PAAD compared with normal pancreatic tissue. However, subsequent analysis of *ATG7* isoform expression in PAAD stages showed similar levels of *ATG7(1)* in all stages and increased *ATG7(2)* expression in Stage IV PAAD (Fig. 1.E), suggesting that *ATG7(2)* could play a role in later stages of PAAD progression, and in metastasis. Additionally, TCGA data of tumours organized by immune subtypes revealed that *ATG7(1)* is more expressed in immunologically quiet cancers while *ATG7(2)* is more expressed in immunologically active cancers (Fig. S1.A), suggesting a role of ATG7(2) in immune modulation. Analysis of survival and new tumour formation in these samples showed that immunologically quiet cancers are significantly linked with greater survival time and higher delay to the apparition of a new tumour compared with immunologically active cancers (Fig. S1.B), suggesting that the effect of ATG7(2) on survival could be the result of immune reprogramming in the tumour microenvironment. Finally, data of tumour infiltrating immune cells was downloaded from the TIMER2.0 database [23], and correlation analysis of *ATG7* expression in these samples revealed that *ATG7* expression is linked with Cancer associated fibroblast (CAFs) and Type-2 macrophage (M2) infiltration in different cancers, including PAAD (Fig. S1.C). This suggests that ATG7 could be involved in the reprogramming of the tumour microenvironment as a result of ATG7(2) activity, thus promoting PAAD progression and increased aggressiveness.

**Figure 1:**
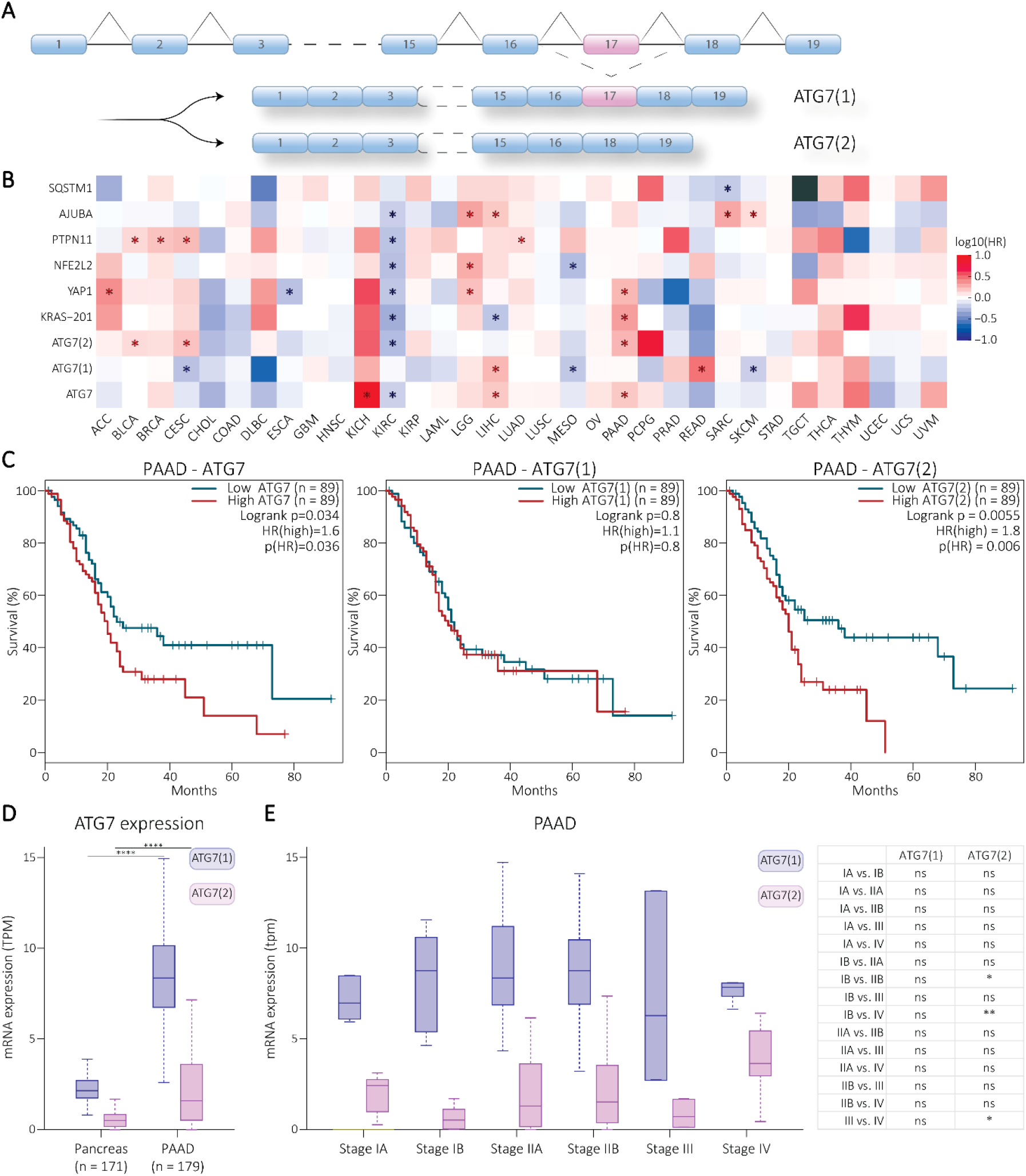
ATG7(2) is linked with poor prognosis in PAAD. **A:** Schematic of the splicing event leading to the transcription of *ATG7(1)* and *ATG7(2)*. **B:** Heatmap showing the correlation between the expression of different mRNA transcripts and survival across cancers (Blue: negative correlation between transcript expression and poor survival; Red: positive correlation between transcript expression and poor survival; *: significant correlation p < 0.05.). **C:** Kaplan-Meier survival plots of *ATG7*; *ATG7(1)* or *ATG7(2)* in PAAD high and low expression groups were determined by median mRNA expression. **D:** Expression plot of *ATG7(1)* and *ATG7(2)* mRNA in PAAD versus pancreas. Multiple t-tests were performed between the Pancreas and PAAD groups, with *: p < 0.05; **: p < 0.01; ***: p < 0.001; ****: p < 0.0001. **E:** Expression plot of *ATG7(1)* and *ATG7(2)* in PAAD clinical stages. One way ANOVA was performed between the different groups, with *: p < 0.05; **: p < 0.01; ***: p < 0.001; ****: p < 0.0001.

### ATG7(2) promotes PAAD cell migration and proliferation *in vitro*

Capan-1 cells were used as an *in vitro* model for PAAD. CRISPR/Cas9 technology was used to selectively knock-out either total ATG7 expression by inducing non-sense mutation in exon 3 of the gene, or ATG7(1) expression by targeting the splice site of exon 17 (Fig. S2.A). In addition, wild-type Capan-1 cells underwent the same CRISPR/Cas9 process in absence of gRNA, and are used and referred to as control cells. Clones harbouring genomic mutations compatible with successful knock-out of ATG7 or ATG7(1) were selected and used for Western-Blotting and qPCR. Both ATG7^−/−^ clones that were obtained, exhibited low levels of *ATG7, ATG7(1)* and *ATG7(2)* mRNA expression compared with the control cells (Fig. S2.B). Western-Blot analysis revealed that neither clone expressed ATG7 protein (Fig. S2.C). Additionally, the mutant cells accumulated p62 protein and lost lipidated LC3B-II compared with the control cells, both indicators of disrupted autophagy activity in the cells. ATG7(1)^−/−^ clones showed no expression of the *ATG7(1)* mRNA, but different levels of *ATG7* mRNA expression, which were reflected by the expression of *ATG7(2)* mRNA (Fig. S2.D). The different levels of *ATG7* expression were also observed at the protein level, and all clones exhibited p62 accumulation and loss of LC3B lipidation compared with the control cells (fig. S2.E). Immunofluorescent staining of the different cell models showed that ATG7 protein is predominantly located in the cytoplasm of the cells (Fig. S2.F). In addition, while control and ATG7(1)^−/−^ cells exhibited similar cytoskeletal architecture, ATG7^−/−^ cells showed rounder shape and fewer filopodia. Together, these results show that ATG7(1) depletion disrupts autophagy activity but does not notably affect global cellular shape, and loss of total ATG7 results in different cellular architecture.

It has been reported that total ATG7 depletion impairs tumour growth in colon adenocarcinoma models [10], and prevents malignancy and metastasis in pancreatic lesions [15], [16], [24]. To assess the effect of ATG7 or ATG7(1) loss in the Capan-1 cells on proliferation and migration, we performed proliferation and wound healing assays using the IncuCyte Live-Cell imaging system. Loss of ATG7(1) led to increased proliferation compared with the control cells, while total ATG7 depletion did not affect the proliferative ability of the cells (Fig. 2.A). Similarly, ATG7(1)^−/−^ cells exhibited higher migration rate than the control cells, while ATG7^−/−^ cells displayed lower migration rate than the control (Fig. 2.B). It is worth noting that the clone with the highest level of ATG7(2) expression was also the clone that exhibited the highest proliferation and migration rates. This suggests that ATG7(1) could be a negative regulator of PAAD cell proliferation and migration, while ATG7(2) seems to function as a positive regulator in a dose-dependent manner. Subsequently, the ATG7(1)^−/−^ clones with the lowest and highest ATG7(2) expression levels, clones 1 and 4 respectively, were selected for further analysis and referred to as ATG7(1)^−/−^; ATG7(2)^low^ (FC = 1.56) and ATG7(1)^−/−^; ATG7(2)^high^ (FC = 7.25). RNA sequencing results confirmed the lack of *ATG7* expression in ATG7^−/−^ cells, as well as the lack of *ATG7(1)* expression in ATG7(1)^−/−^ cells (Figure S3.A). The results also confirmed that ATG7(1)^−/−^ clone 1 expresses similar *ATG7(2)* levels as the control cells, while ATG7(1)^−/−^ clone 4 expresses higher *ATG7(2)* levels. This indicates that ATG7(1) depletion is sufficient to block autophagy activity without affecting the function of other ATG7 isoforms. Analysis of RNA sequencing data also revealed altered gene expression profiles between the different conditions (Fig. S3.B-C), while having similar sequencing profiles (Fig S3.D), confirming the robustness of the analysis. This shows that the differences observed between the different conditions are the result of the gene expression profiles, not differences in their sequencing analysis.

**Figure 2:**
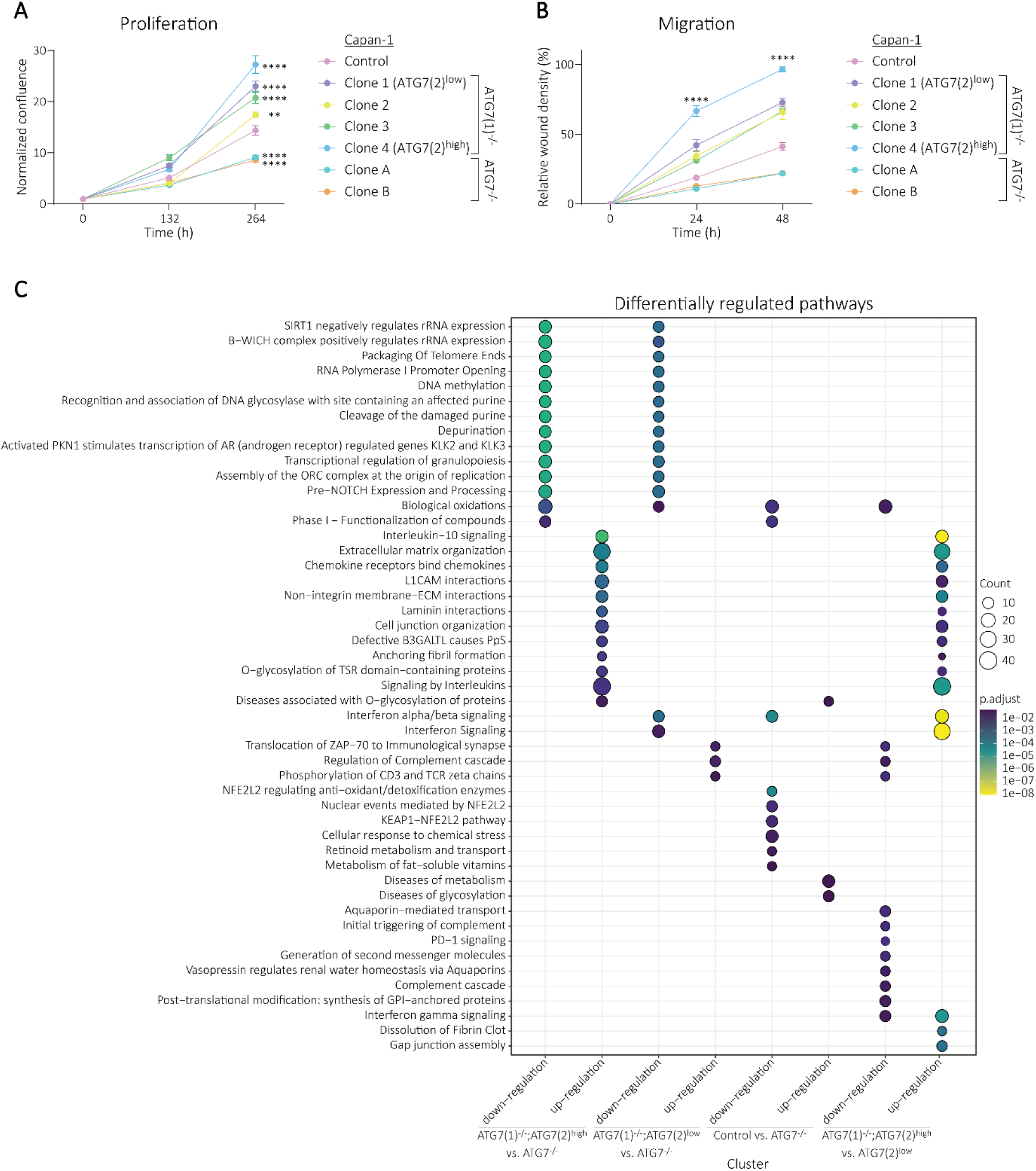
ATG7(2) drives PAAD cell proliferation and migration, and regulates the expression of genes involved in ECM organization, immune signalling and cell-cell interaction. **A:** Normalized proliferation ratios of Capan-1 control, ATG7(1)^−/−^ and ATG7^−/−^ cells at 132h and 264h timepoints after seeding. **B:** Normalized migration ratios of Capan-1 control, ATG7(1)^−/−^ and ATG7^−/−^ cells at 24h and 48h timepoints after wound making. **A-B:** Two way ANOVA was performed between the control and every other condition, with *: p < 0.05; **: p < 0.01; ***: p < 0.001; ****: p < 0.0001. **C:** Functional enrichment of response genes (p < 0.05; 2-fold change) between Capan-1 control, ATG7(1)^−/−^; ATG7(2)^high^, ATG7(1)^−/−^; ATG7(2)^low^ and ATG7^−/−^ cells using Reactome_2022, from RNA sequencing.

Control cells were cultivated in full medium, or under glucose starvation for 4 hours, and levels of *ATG7, ATG7(1)*, and *ATG7(2)* mRNA expression were measured using qPCR. This showed that *ATG7* and *ATG7(2)* expression was induced by nutrient stress, while *ATG7(1)* expression did not change significantly (Fig. S3.E). This implies that ATG7(2) could be involved in stress-coping mechanisms. RNA sequencing analysis also revealed that ATG7(1)^−/−^; ATG7(2)^high^ cells expressed the highest levels of the proliferation marker *Ki67*, which was consistent with the results from the proliferation assay (Fig. S3.F). Gene enrichment analysis of the RNA sequencing data showed down-regulation of metabolic pathways and NRF2 signalling in control cells compared with ATG7^−/−^ cells (Fig. 2.C, Table S1). Both ATG7(1)^−/−^; ATG7(2)^high^ and ATG7(1)^−/−^; ATG7(2)^low^ cells showed down-regulated DNA replication and transcription. ATG7(1)^−/−^; ATG7(2)^high^ cells exhibited up-regulation of genes involved in ECM organization, cell-cell interaction and interleukin signalling compared with ATG7(1)^−/−^; ATG7(2)^low^ and ATG7^−/−^ cells. The expression of genes involved in immune pathways such as ZAP-70 translocation and CD3 phosphorylation were up-regulated in ATG7(1)^−/−^; ATG7(2)^low^ cells compared with ATG7(1)^−/−^; ATG7(2)^high^ and ATG7^−/−^ cells. In addition, ATG7(1)^−/−^; ATG7(2)^high^ cells exhibited higher expression of the *CX3CL1* and *CXCL11* mRNAs, encoding cytokines which promote immune evasion in PAAD [25], [26], as well as higher expression of RNAs encoding major histocompatibility complex I (MHC-I) components (Fig. S3.G). Together, these results suggest that ATG7(2) promotes PAAD progression *in vitro*, down-regulates DNA replication and transcription in a dose-independent manner. Moreover, high levels of ATG7(2) promote ECM organization, cell-cell interaction and interleukins signalling, and that lower levels of ATG7(2) positively regulate other immune pathways, indicating a potential dose-dependent role of ATG7(2) in PAAD.

### ATG7(2) regulates chemokine expression and secretion

To further explore the effect of ATG7(2) on tumour immunoactivity, we performed targeted analysis of the RNA-seq data to evaluate the mRNA expression of different cytokines involved in the recruitment and polarization of tumour associated macrophages (TAMs), myeloid-derived suppressor cells (MDSCs), and regulatory T cells (Treg). Luminex multiplex quantification assay was used to measure protein levels in the cell culture supernatant, which was normalized to total protein amount from the cells to quantify secreted cytokine amount. Results showed that ATG7(1)^−/−^; ATG7(2)^high^ cells consistently expressed significantly higher mRNA levels of *CCL2, CXCL10, CCL5, IL-6* and *IL-8* than the control, ATG7^−/−^ and ATG7(1)^−/−^; ATG7(2)^low^ cells (Fig. 3.A). However, both ATG7(1)^−/−^; ATG7(2)^low^ and ATG7(2)^high^cell lines secreted greater amounts of these cytokines than the control and ATG7^−/−^ cells, with ATG7(1)^−/−^; ATG7(2)^low^ cells secreting higher amounts of CCL2, IL-6 and IL-8 than ATG7(1)^−/−^; ATG7(2)^high^ cells (Fig. 3.B). ATG7(1)^−/−^; ATG7(2)^high^ cells also exhibited higher levels of *CCL3, CXCL1, Galectin-1, IL-1α* and *GM-CSF* mRNA than the control, ATG7^−/−^ and ATG7(1)^−/−^; ATG7(2)^low^ cells (Fig. 3.C). While this trend was conserved at the secreted protein level for CCL3 and Galectin-1, ATG7(1)^−/−^; ATG7(2)^high^ cells displayed similar secretion levels of CXCL1 as the control cells, and ATG7(1)^−/−^; ATG7(2)^low^ cells exhibited higher secreted levels of CXCL1, IL-1α and GM-CSF than the control cells (Fig. 3.D). These results suggest that ATG7(2) activity not only affects cytokine expression, but also their secretion.

**Figure 3:**
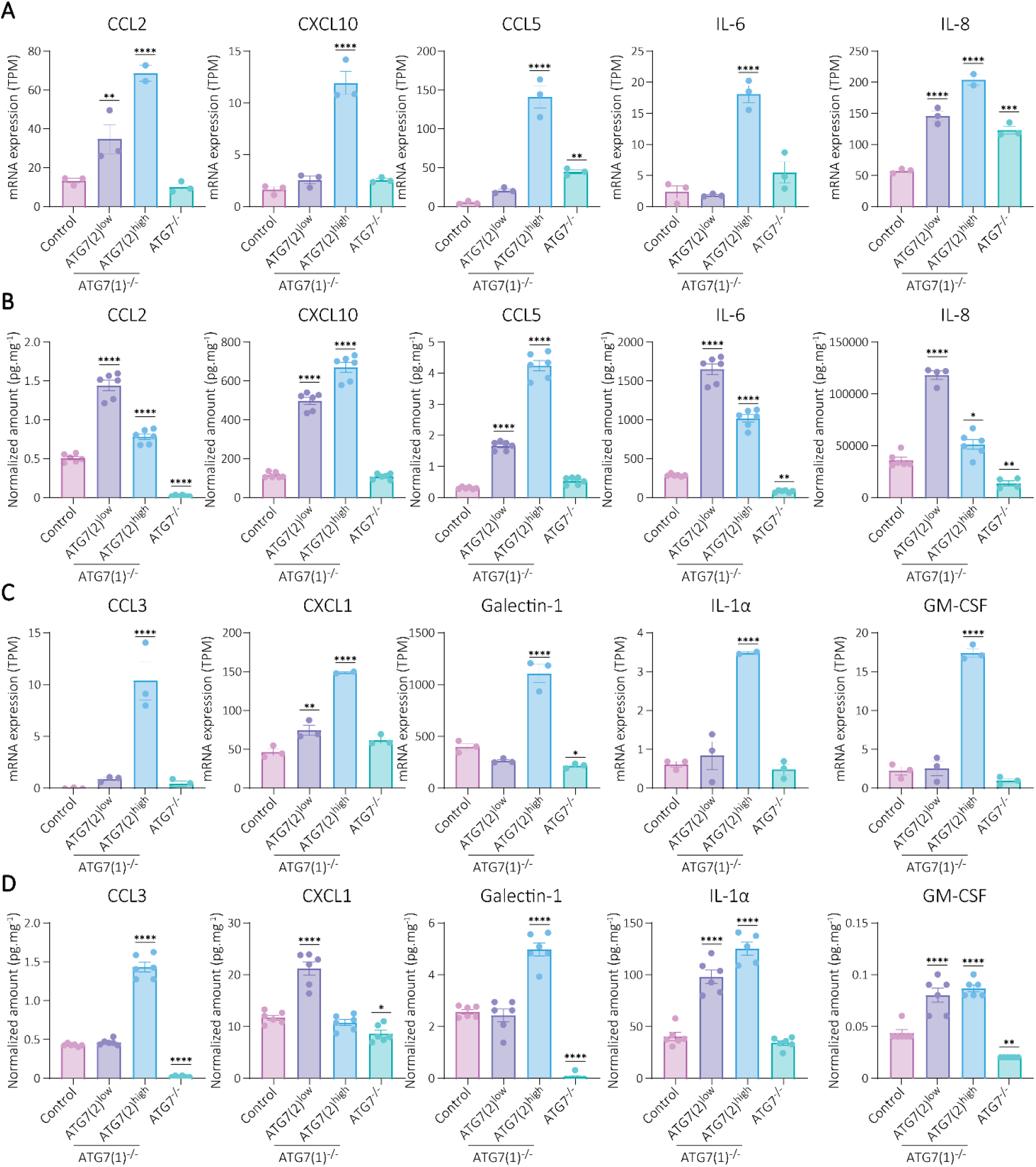
ATG7(2) regulates the expression and secretion of cytokines and chemokines. **A:** mRNA expression of factors involved in Treg, TAMs and NK cells recruitment and polarization, from RNA sequencing. **B:** Protein secretion of factors involved in Treg, TAMs and NK cells recruitment and polarization, from Multiplex Luminex® quantification. **C:** mRNA expression of factors involved in T cells, Neutrophils and MDSCs recruitment and polarization, from RNA sequencing. **D:** Protein secretion of factors involved in T cells, Neutrophils and MDSCs recruitment and polarization, from Multiplex Luminex® quantification. **A-D:** One way ANOVA was performed between the control and every other condition, with *: p < 0.05; **: p < 0.01; ***: p < 0.001; ****: p < 0.0001.

Tumour immunity and ECM organization have been linked to the Hippo pathway in numerous studies [27], [28]. The Hippo pathway promotes YAP phosphorylation and degradation, preventing it from entering the nucleus where, along with SHP2, it binds to the transcription factor TEAD to induce the expression of genes involved in cell proliferation, ECM organization and immune regulation [27], [28], [29]. This pathway is known to be frequently dysregulated in cancer [30], [31]. To assess the potential effect of ATG7(2) on Hippo regulation and YAP nuclear activity, we performed targeted analysis of the RNA-seq data to evaluate the mRNA expression of different genes, the expression of which is promoted by YAP. This revealed that ATG7(1)^−/−^; ATG7(2)^high^ cells consistently expressed significantly more *YAP1* mRNA and of its target genes *AREG, AXL, CCND1* and *CYR61*, compared with the control cells (Fig. S4.A). This suggests that ATG7(2) negatively regulates the Hippo pathway. Interestingly, we have previously observed that ATG7(2) interacts with the LIM protein Ajuba [32], a known negative regulator of the Hippo pathway. To further explore the effect of ATG7(2) on YAP activity, we performed subcellular fractionation and evaluated cytoplasmic and nuclear levels of Ajuba, SHP2 and YAP. We observed that there was no ATG7 protein in the nuclear fraction of either control or ATG7(1)^−/−^; ATG7(2)^high^ cells (Fig. S4.B). Furthermore, the results showed accumulation of SHP2, Ajuba and YAP protein in ATG7^−/−^ cells compared with the control, as previously reported for YAP [33]; ATG7(1)^−/−^; ATG7(2)^high^ cells showed even more accumulation. Quantification of cytoplasmic and nuclear protein amounts showed decreased Ajuba nuclear translocation in both ATG7^−/−^ and ATG7(1)^−/−^; ATG7(2)^high^ cells compared with the control, decreased SHP2 nuclear translocation in ATG7^−/−^ cells compared with control and ATG7(1)^−/−^; ATG7(2)^high^ cells, and increased YAP nuclear translocation in ATG7(1)^−/−^; ATG7(2)^high^ cells compared with control and ATG7^−/−^ cells (Fig. S4.C). This was also observed when staining the control, ATG7(1)^−/−^ and ATG7^−/−^ cells for YAP and imaging them using confocal microscopy (Fig. S4.D). Together, these results show that both ATG7(1) and ATG7(2) are active in the cytoplasm but not in the nucleus and suggests that ATG7(1) could be involved in YAP degradation through autophagy while ATG7(2) might prevent YAP degradation in an autophagy independent manner.

### ATG7(2) inhibition decreases PAAD cell migration and proliferation *in vitro*

To study the therapeutic potential of targeting ATG7 isoforms in PAAD, we designed siRNA sequences targeting *ATG7, ATG7(1)* or *ATG7(2)* specifically (Fig. S5.A). The specificity of the designs was confirmed in Atg7^−/−^ mouse embryonic fibroblasts (MEFs) overexpressing either ATG7(1) or ATG7(2) (Fig. S5.B-C). The efficiency of the siATG7(2) designs was assessed in Capan-1 control and ATG7(1)^−/−^; ATG7(2)^high^ cells, and the most efficient design was selected (Fig. S5.D-E).

Proliferation and migration assays were performed on control, ATG7(1)^−/−^; ATG7(2)^low^ and ATG7(1)^−/−^; ATG7(2)^high^ cells, transfected with control siRNA or siATG7(2) every 72 hours. Both ATG7(1)^−/−^ cell lines displayed substantially slower proliferation rates, and all three cell lines displayed significantly slower migration rates upon siATG7(2) treatment compared with the control siRNA (Fig. 4.A-B). RNA sequencing was performed on Panc 08.13 and Capan-1 control, ATG7(1)^−/−^; ATG7(2)^low^ and ATG7(1)^−/−^; ATG7(2)^high^ cells, treated with control siRNA or siATG7(2) for 72 hours. The pancreatic cancer cells Panc 08.13 were used in addition to the Capan-1 cells to compare the effect in cells with different background and genome, thus assessing the context-dependent effect of the phenotype. This revealed that knocking-down ATG7(2) significantly decreased *ATG7* mRNA expression in Capan-1 ATG7(1)^−/−^; ATG7(2)^low^ (p = 0.023) and ATG7(1)^−/−^; ATG7(2)^high^ (p = 0.0007) cells, and to a lesser extent in control cells (Fig. S6.A). Additionally, our data showed limited gene expression differences between the siATG7(2) and control condition in each cell line (Fig. S6.B-C). Interestingly, enrichment analysis of the RNA sequencing data revealed that ATG7(2) inhibition affects the expression of a limited number of genes (31 – 182), but with important involvement in immune response and immune cell chemotaxis, in Capan-1 control and ATG7(1)^−/−^; ATG7(2)^high^ cells (Fig. 4.C, Table S2). Interestingly, *HMGA2* and *CLDN1* appeared amongst the five most differentially regulated genes upon ATG7(2) inhibition in Capan-1 control, ATG7(1)^−/−^; ATG7(2)^high^ and ATG7(1)^−/−^; ATG7(2)^low^ (Fig. 4.D, Fig. S6.D, Table S3). While the expression of *HMGA2* and *CLDN1* were significantly up-regulated, *CYB5R4* expression was significantly down-regulated upon ATG7(2) inhibition in all three cell lines (Fig. S6.E). In addition, ATG7(2) inhibition also affected the transcription of *CCL2, CXCL10, CCL5, IL-6* and *IL-8* (Fig. 4.E), and their secretion, with the exception of IL-8 secretion (Fig. 4.F).

**Figure 4:**
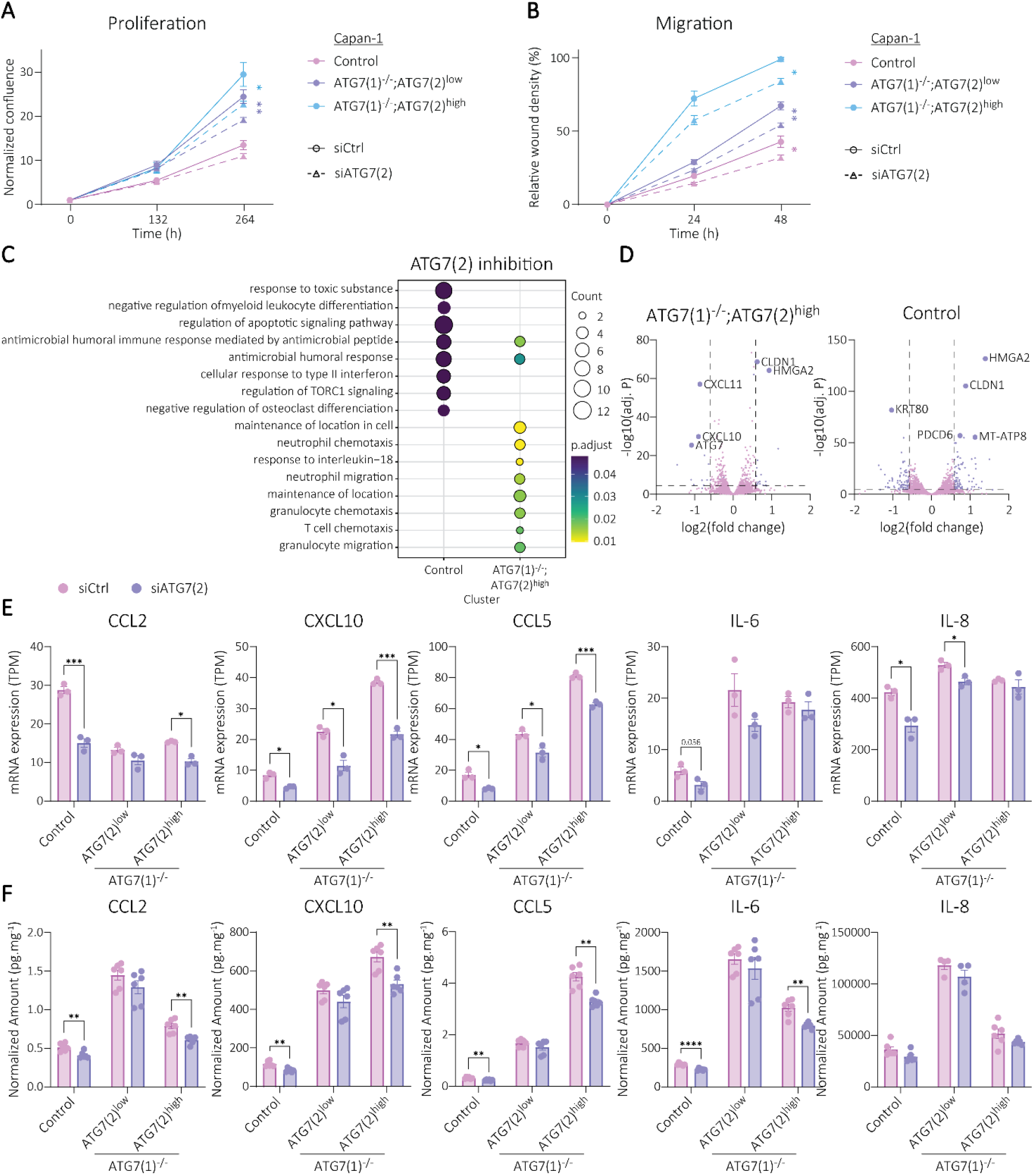
ATG7(2) inhibition reduces PAAD cell migration and proliferation *in vitro*, and affects immune signalling. **A:** Normalized proliferation ratios of Capan-1 control and ATG7(1)^−/−^ cells after siATG7(2) or control siRNA treatment every 72h, at 132h and 264h timepoints after seeding. **B:** Normalized migration ratios of Capan-1 control and ATG7(1)^−/−^ cells after siATG7(2) or control siRNA treatment every 72h, at 24h and 48h timepoints after wound making. **C:** Functional enrichment of significantly differentially regulated genes (p < 0.05; 1.5-fold change) between control siRNA and siATG7(2) in Capan-1 Control and ATG7(1)^−/−^; ATG7(2)^high^ cells using Reactome_2022, from RNA sequencing. **D:** Volcano plots depicting differentially expressed genes in Control and ATG7(1)^−/−^; ATG7(2)^high^ cells when treated with siATG7(2) for 72h. Pink dots represent genes that are not differentially expressed, while purple dots represent genes that are differentially expressed. The 5 genes with the lowest adjusted p value are labelled. **E:** mRNA expression of factors involved in Treg, TAMs and NK cells recruitment and polarization, after siATG7(2) or control siRNA treatment for 72h. From RNA sequencing. **F:** Protein secretion of factors involved in Treg, TAMs and NK cells recruitment and polarization, after siATG7(2) or control siRNA treatment for 72h. From Multiplex Luminex® quantification. **A-B**,**E-F:** Multiple t-tests were performed between the control and the siATG7(2) condition, with *: p < 0.05; **: p < 0.01; ***: p < 0.001; ****: p < 0.0001.

The expression of *CXCL11* was down-regulated upon ATG7(2) inhibition, but not *CX3CL1* expression, and not all MHC-I components were reduced in expression upon ATG7(2) inhibition (Fig. S6.F). In Panc 08.13 cells, ATG7(2) inhibition significantly reduced the secretion of IL-6, CXCL10, IL-1α, CCL5, CCL3 and GM-CSF (Fig. S7.A-B). In Capan-1 cells, ATG7(2) inhibition significantly decreased the expression of *CXCL1, IL-1α* and *GM-CSF* (Fig. S8.A), and the secretion of CCL3, CXCL1, IL-1α and GM-CSF (Fig. S8.B), although the effect of ATG7(2) inhibition on the expression and secretion of these cytokines varied between conditions. Similar to ATG7(2) inhibition, YAP1 inhibition, performed with siYAP1 or control siRNA transfection every 72h, led to decreased proliferation and migration rates compared with the control, in Capan-1 control, ATG7(1)^−/−^; ATG7(2)^low^, ATG7(1)^−/−^; ATG7(2)^high^, ATG7^−/−^ cells (Fig. S8.C-D), and Panc 08.13 cells (Fig. S8.E-F). To assess the impact of ATG7(2) knock-down on YAP transcriptional activity, we analysed the expression of genes positively regulated by nuclear YAP activity in our RNA sequencing data. This revealed that ATG7(2) inhibition had no significant effect on the expression of *YAP1, AREG, AXL, CCND1, CYR61* and *LATS2* in any of the Capan-1 conditions (Fig. S9.A), suggesting that ATG7(2) inhibition does not affect YAP transcriptional activity. Together, these results suggest that ATG7(2) is involved in the regulation of tumour-protective cytokine expression and secretion, independently of YAP activity.

### ATG7(2) inhibition decreases PAAD progression *in vivo*

We used BALB/c nude mice to transplant Capan-1 control, ATG7(1)^−/−^; ATG7(2)^low^, ATG7(1)^−/−^; ATG7(2)^high^ and ATG7^−/−^ cells and study their progression *in vivo*. BALB/c nude mice are immunocompromised due to lack of thymus, and therefore T cell populations as well. All four control groups were treated twice weekly with control siRNA via intratumoral injections, and two additional control and ATG7(1)^−/−^; ATG7(2)^high^ groups that were treated twice weekly with siATG7(2) (Fig. 5.A). Whereas the weight of the animals was not affected by the treatment (Fig. S10.A), control tumours progressed significantly faster than ATG7(1)^−/−^ and ATG7^−/−^ tumours, with the ATG7^−/−^ group exhibiting the slowest progression rate, and ATG7(1)^−/−^; ATG7(2)^low^ tumours progressing significantly faster than ATG7(1)^−/−^; ATG7(2)^high^ tumours (Fig. 5.B). This shows that ATG7(2) supports PAAD tumour growth, though to a lesser extent than *in vitro*.

**Figure 5:**
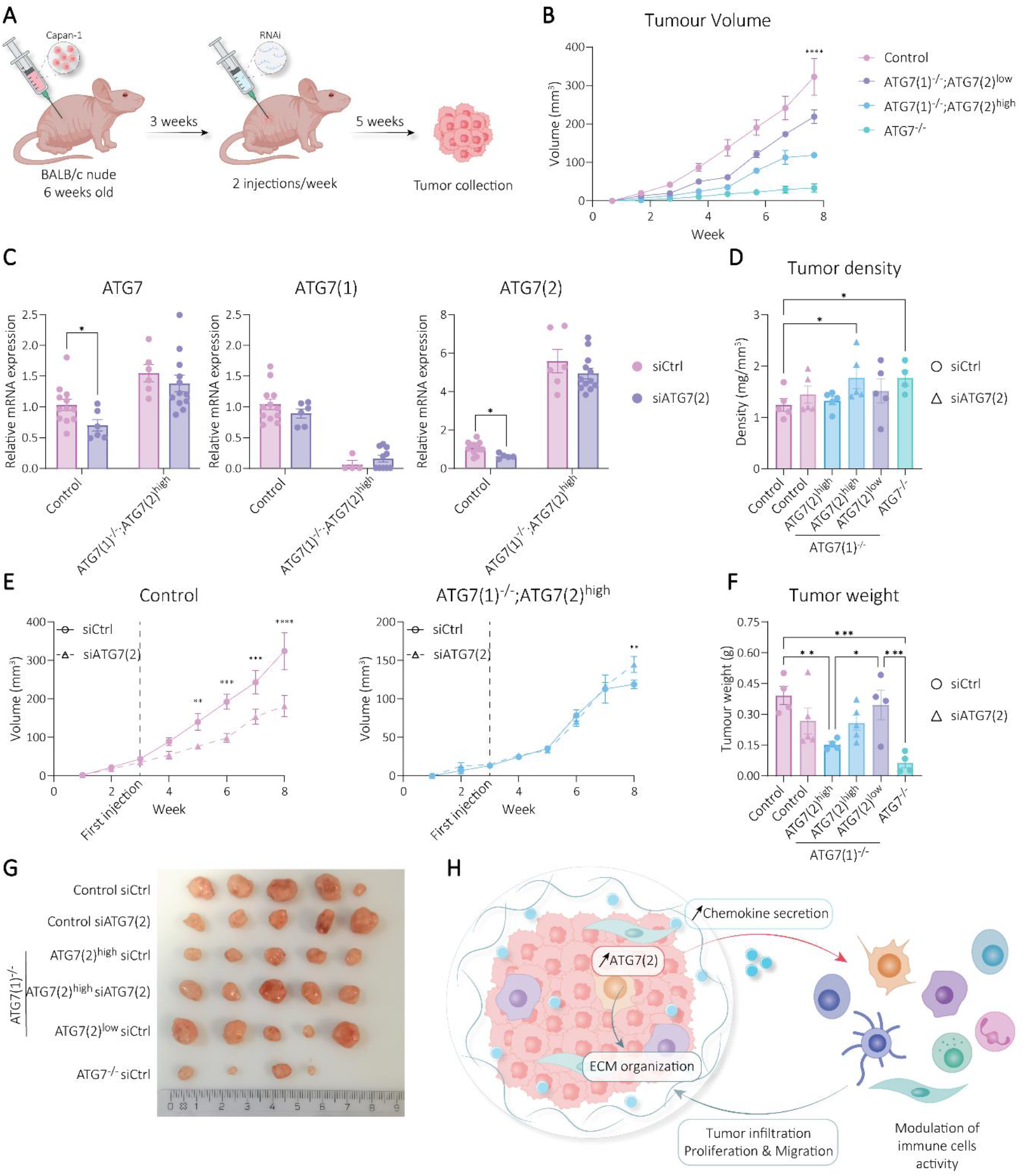
ATG7(2) contributes to PAAD progression *in vivo*. **A:** Schematic of the xenograft study. Capan-1 control, ATG7(1)^−/−^; ATG7(2)^high^, ATG7(1)^−/−^; ATG7(2)^low^ and ATG7^−/−^ cells were injected subcutaneously in the right flank of BALB/c nude mice. Capan-1 control and ATG7(1)^−/−^; ATG7(2)^high^ tumours were treated with siATG7(2) or control siRNA, while ATG7(1)^−/−^; ATG7(2)^low^ and ATG7^−/−^ tumours were treated with control siRNA only. **B:** Evolution of the tumor volume in each group treated with control siRNA (n=5). **C:** mRNA expression of *ATG7, ATG7(1), ATG7(2)* in Capan-1 Control and ATG7(1)^−/−^; ATG7(2)^high^ cohorts treated with control siRNA or siATG7(2), quantified with qPCR at the end of the study. **D:** Tumour densities at the end of the study. **E:** Evolution of the tumour volume in Capan-1 control and ATG7(1)^−/−^; ATG7(2)^high^ cohorts treated with control siRNA or siATG7(2) (n=5). **F:** Tumour weights at the end of the study. **B:** One way ANOVA was performed between the control and every other condition / **C-F:** Multiple t-tests were performed between the control and the siATG7(2) condition, with *: p < 0.05; **: p < 0.01; ***: p < 0.001; ****: p < 0.0001. **G:** Image of the tumours at the end of the study. **H:** Graphical model of the role of ATG7(2) in promoting PAAD progression through immune modulation.

Inhibition of ATG7(2) by siRNA was statistically significant at mRNA level in the control group, but not in the ATG7(1)^−/−^; ATG7(2)^high^ group (Fig. 5.C). This lack of efficiency in the ATG7(1)^−/−^; ATG7(2)^high^ group could be explained by a dense extracellular-matrix, as suggested by the previous RNA sequencing results, as well as a significantly higher tumour density than the control group (Fig. 5.D). While ATG7(2) inhibition was unsuccessful in the ATG7(1)^−/−^; ATG7(2)^high^ group, it successfully and substantially slowed down tumour progression from the second week of treatment in the Control group (Fig. 5.E). At the end of the study, the tumours were collected, measured, weighed and photographed, and the postmortem tumour weight and volume were consistent with the differences in tumour progression observed between the different groups (Fig. 5.F-G, Fig. S10.B-C). Together, these results suggest that ATG7(2) activity is involved in the modulation of the tumour micro-environment to promote pancreatic cancer progression and immune evasion (Fig. 5.H). This brings light to the promising potential of ATG7(2) as a therapeutic target for the treatment of PAAD, which could be used in combination with immunotherapies.

## DISCUSSION

While ATG7 is best known for its role in autophagy and cellular homeostasis, increasing evidence indicates that it also has autophagy-independent functions that contribute to tumour progression. In pancreatic cancer (PAAD), our findings identify the short isoform ATG7(2), which lacks canonical autophagy activity, as a key driver of this tumour effect. Analysis of publicly available datasets showed that high *ATG7(2)* expression is associated with poor prognosis in PAAD, whereas expression of the canonical isoform *ATG7(1)* is not. Our analyses further suggest a link between ATG7(2) and immune modulation across cancer types. Using isoform-specific knock-out models, we demonstrate that ATG7(2) promotes PAAD cell proliferation and migration *in vitro*. Transcriptomic analysis further revealed that ATG7(2) regulates pathways related to immune signalling, extracellular matrix organisation, and cell–cell interactions in a dose-dependent manner. In addition, ATG7(2) influenced both cytokine expression and secretion, indicating a role in shaping the tumour microenvironment, while its effects appeared to be partially counteracted by the presence of ATG7(1). Interestingly, the analysis of RNA sequencing data revealed that macro autophagy was not significantly dysregulated in ATG7^−/−^ cells compared with control cells. Though the cells showed clear p62 accumulation and absence of atg8ylation, it is possible that the cells adapted to ATG7 loss by rewiring toward ATG5/ATG7-independent autophagy [34].

Through autophagy, ATG7 has been linked to innate immunity [35], and indeed, ATG7 has been implicated in the activation of LC3-associated phagocytosis (LAP), an antimicrobial pathway at the crossroad between phagocytosis and autophagy [36]. This process is used by macrophages to degrade pathogens, and LAP is believed to attenuate the inflammation caused by dying cells [37]. Atg7 promotes melanoma progression *in vivo* by preventing myelosuppressive immune cell infiltration in the tumour microenvironment [7]. Another study showed that Atg7 inhibition improved angiogenesis and tumour suppressive cell infiltration in the microenvironment of several solid tumours [38]. Interestingly, it has been shown that ATG7 regulates angiogenesis independently of autophagy, suggesting that the effect observed in that study [38] could be autophagy-independent [39]. In the same study, results showed that combination of Atg7 inhibition and PD-1 inhibition significantly reduced tumour growth compared with PD-1 inhibition alone [38]. Together, these findings support a broader role for ATG7 in regulating tumour immunity and the microenvironment, consistent with our observations that ATG7(2) modulates cytokine expression and immune-related pathways in PAAD cells.

Given the role of ATG7(2) in modulating cytokine expression and immune-related pathways, our findings suggest that it may also influence tumour immune evasion mechanisms. Immune checkpoint proteins, which are expressed on the surface of cancer cells, enable tumours to evade immune detection and promote the polarization of immune cells toward tumour-supportive phenotypes, such as tumour-associated macrophages (TAMs) [40]. Inhibition of these checkpoints restores immune recognition and limits tumour progression, forming the basis of widely used immunotherapies [41], [42]. Given the current findings, targeting ATG7(2) may represent a complementary strategy to enhance anti-tumour immune responses. Our RNA-seq data indicating a role for ATG7(2) in immune modulation support the potential of combining ATG7(2) inhibition with checkpoint blockade therapies, such as PD-1/PD-L1 inhibitors. Consistent with this, previous studies have shown that targeting ATG7 or autophagy pathways can improve the efficacy of anti-PD-1 treatment [9], [38].

Interestingly, despite having the highest proliferation and migration rates i*n vitro*, ATG7(1)^−/−^; ATG7(2)^high^ cells did not lead to increased tumour progression when compared with control cells *in vivo*. Similarly, ATG7(1)^−/−^; ATG7(2)^low^ cells displayed enhanced proliferation compared to control cells *in vitro*, but not *in vivo*. This contrast highlights the potential importance of the tumour microenvironment in modulating the effects of ATG7(2), suggesting that its pro-tumorigenic role is not entirely cell-intrinsic but likely context-dependent. This difference could be explained by the differential cytokine expression and secretion in addition to the internal proliferative and migrative effect of ATG7(2) expression in PAAD cells. Indeed, ATG7(1)^−/−^, and especially ATG7(1)^−/−^; ATG7(2)^high^ cells, expressed and secreted higher levels of numerous cytokines compared to control cells. As ATG7 has previously been linked to CD8+ intratumoral infiltration [7], [8], it is possible that the lack of T cells in BALB/c nude mice impaired the potential of ATG7(1)^−/−^ cells *in vivo*.

Finally, we assessed the therapeutic potential of ATG7(2) inhibition using siRNA. Our results showed that ATG7(2) inhibition decreased cell migration and proliferation *in vitro*, as well as the expression and secretion of various cytokines, suggesting that ATG7(2) is involved in the modulation of the tumour microenvironment through ECM remodelling, and immune cell modulation. Importantly, ATG7(2) inhibition also significantly slowed tumour progression *in vivo*, highlighting its potential as a therapeutic target in PAAD. However, siRNA-mediated knockdown was ineffective in ATG7(1)-/-; ATG7(2)high tumours. One possibility is that this was due to increased tumour density and extracellular matrix deposition, which are known to limit oligonucleotide delivery [43], [44]. This limitation underscores a key challenge for therapeutic targeting of ATG7(2), but also points to potential strategies to enhance treatment efficacy, such as combining ATG7(2) inhibition with approaches that improve tumour penetration, including matrix remodelling or VEGF inhibition [45], [46].

The role of ATG7 has been shown to be context dependent, with p53 mutational status being reported to be affected by, and affect ATG7 function in PAAD [15], [16], [24], [47]. In addition, ATG7 encoded by a plasmid containing *ATG7(2)* cDNA, has been reported to directly interact with p53 [48]. The role of ATG7(2) on immune signalling and the context-dependent role of ATG7 in PAAD call for a deeper analysis of ATG7(2) function in an immunocompetent PAAD murine model, with different Trp53 status.

Together, these findings identify ATG7(2) as a driver of pancreatic cancer progression through autophagy-independent mechanisms and highlight its potential as a therapeutic target in PAAD.

## ACKNOWLEDGEMENTS

This project was supported by the Icelandic Research Fund grant 228586-051. We thank Bylgja Hilmarsdottir for providing us with the Capan-1 cells and Jaclyn Long for providing us with the Panc 08.13 cells. We thank Siggeir Fannar Brynjolfsson for his valuable guidance on the Luminex assays, as well as Bergthora Eiriksdottir and Thora Jona Dagbjartsdottir for their help and guidance in the animal experiments. We are also very grateful to Eirikur Steingrimsson, Paula Fernandez Palanca and Kevin Ryan for revising this manuscript.

## AUTHOR CONTRIBUTIONS

M. Ögmundsdóttir and C. Larat conceived the project and designed the study. M. Ogmundsdottir supervised the project. C. Larat analysed the clinical data, performed the experiments, designed siRNAs and primers, and wrote the manuscript. A. L. G. de Lomana supervised and provided code for the RNA sequencing analysis, and V. Hjaltalin designed the CRISPR strategy. M. Ogmundsdottir, A. L. G. de Lomana and V. Hjaltalin critically revised the manuscript. All authors have read and agreed to the published version of the manuscript.

## REFERENCES

[1] Cao, J. Li, K. Yang, and D. Cao, ‘An overview of autophagy: Mechanism, regulation and research progress’, Bull. Cancer (Paris), vol. 108, no. 3, pp. 304–322, Mar. 2021, doi: 10.1016/j.bulcan.2020.11.004.

[2] N. S. Vargas, M. Hamasaki, T. Kawabata, R. J. Youle, and T. Yoshimori, ‘The mechanisms and roles of selective autophagy in mammals’, Nat. Rev. Mol. Cell Biol., vol. 24, no. 3, pp. 167–185, Mar. 2023, doi: 10.1038/s41580-022-00542-2.

[3] R. Miller and A. Thorburn, ‘Autophagy and organelle homeostasis in cancer’, Dev. Cell, vol. 56, no. 7, pp. 906–918, Apr. 2021, doi: 10.1016/j.devcel.2021.02.010.

[4] W. Li et al., ‘Selective autophagy of intracellular organelles: recent research advances’, Theranostics, vol. 11, no. 1, pp. 222–256, 2021, doi: 10.7150/thno.49860.

[5] Devis-Jauregui, N. Eritja, M. L. Davis, X. Matias-Guiu, and D. Llobet-Navàs, ‘Autophagy in the physiological endometrium and cancer’, Autophagy, vol. 17, no. 5, pp. 1077–1095, May 2021, doi: 10.1080/15548627.2020.1752548.

[6] W. Zhang et al., ‘Autophagic Schwann cells promote perineural invasion mediated by the NGF/ATG7 paracrine pathway in pancreatic cancer’, J. Exp. Clin. Cancer Res. CR, vol. 41, no. 1, p. 48, Feb. 2022, doi: 10.1186/s13046-021-02198-w.

[7] M. Zimmerman, E. Suh, S. R. Smith, and G. P. Souroullas, ‘Stat3-mediated Atg7 expression regulates anti-tumor immunity in mouse melanoma’, Cancer Immunol. Immunother., vol. 73, no. 11, p. 218, Sep. 2024, doi: 10.1007/s00262-024-03804-4.

[8] M. Zhu et al., ‘ATG7 is associated with immune infiltration and affects the prognosis of colorectal cancer patients’, Asian J. Surg., vol. 49, no. 2, pp. 700–712, Feb. 2026, doi: 10.1016/j.asjsur.2025.08.234.

[9] W. Zhang et al., ‘Inhibition of autophagy-related protein 7 enhances anti-tumor immune response and improves efficacy of immune checkpoint blockade in microsatellite instability colorectal cancer’, J. Exp. Clin. Cancer Res. CR, vol. 43, p. 114, Apr. 2024, doi: 10.1186/s13046-024-03023-w.

[10] D. Arensman et al., ‘Anti-tumor immunity influences cancer cell reliance upon ATG7’, Oncoimmunology, vol. 9, no. 1, p. 1800162, doi: 10.1080/2162402X.2020.1800162.

[11] Rui, L. Zhou, and S. He, ‘Cancer immunotherapies: advances and bottlenecks’, Front. Immunol., vol. 14, p. 1212476, 2023, doi: 10.3389/fimmu.2023.1212476.

[12] S. O’Donnell, M. W. L. Teng, and M. J. Smyth, ‘Cancer immunoediting and resistance to T cell-based immunotherapy’, Nat. Rev. Clin. Oncol., vol. 16, no. 3, pp. 151–167, Mar. 2019, doi: 10.1038/s41571-018-0142-8.

[13] Vasan, J. Baselga, and D. M. Hyman, ‘A view on drug resistance in cancer’, Nature, vol. 575, no. 7782, pp. 299–309, Nov. 2019, doi: 10.1038/s41586-019-1730-1.

[14] M. Zhu et al., ‘ATG7 is associated with immune infiltration and affects the prognosis of colorectal cancer patients’, Asian J. Surg., vol. 49, no. 2, pp. 700–712, Feb. 2026, doi: 10.1016/j.asjsur.2025.08.234.

[15] A. Yang et al., ‘Autophagy is critical for pancreatic tumor growth and progression in tumors with p53 alterations’, Cancer Discov., vol. 4, no. 8, pp. 905–913, Aug. 2014, doi: 10.1158/2159-8290.CD-14-0362.

[16] S. Long et al., ‘ATG7 is a haploinsufficient repressor of tumor progression and promoter of metastasis’, Proc. Natl. Acad. Sci. U. S. A., vol. 119, no. 28, p. e2113465119, Jul. 2022, doi: 10.1073/pnas.2113465119.

[17] H. Ogmundsdottir et al., ‘A short isoform of ATG7 fails to lipidate LC3/GABARAP’, Sci. Rep., vol. 8, no. 1, p. 14391, Sep. 2018, doi: 10.1038/s41598-018-32694-7.

[18] M.H. Ostacolo et al., ‘ATG7(2) Interacts With Metabolic Proteins and Regulates Central Energy Metabolism’, Traffic Cph. Den., vol. 25, no. 4, p. e12933, Apr. 2024, doi: 10.1111/tra.12933.

[19] Chen, Y. Zhou, Y. Chen, and J. Gu, ‘fastp: an ultra-fast all-in-one FASTQ preprocessor’, Bioinformatics, vol. 34, no. 17, pp. i884–i890, Sep. 2018, doi: 10.1093/bioinformatics/bty560.

[20] L. Bray, H. Pimentel, P. Melsted, and L. Pachter, ‘Near-optimal probabilistic RNA-seq quantification’, Nat. Biotechnol., vol. 34, no. 5, pp. 525–527, May 2016, doi: 10.1038/nbt.3519.

[21] I. Love, W. Huber, and S. Anders, ‘Moderated estimation of fold change and dispersion for RNA-seq data with DESeq2’, Genome Biol., vol. 15, no. 12, p. 550, 2014, doi: 10.1186/s13059-014-0550-8.

[22] Tang, B. Kang, C. Li, T. Chen, and Z. Zhang, ‘GEPIA2: an enhanced web server for large-scale expression profiling and interactive analysis’, Nucleic Acids Res., vol. 47, no. W1, pp. W556–W560, Jul. 2019, doi: 10.1093/nar/gkz430.

[23] T. Li et al., ‘TIMER2.0 for analysis of tumor-infiltrating immune cells’, Nucleic Acids Res., vol. 48, no. W1, pp. W509–W514, Jul. 2020, doi: 10.1093/nar/gkaa407.

[24] T. Rosenfeldt et al., ‘p53 status determines the role of autophagy in pancreatic tumour development’, Nature, vol. 504, no. 7479, pp. 296–300, Dec. 2013, doi: 10.1038/nature12865.

[25] A. Chaudhri et al., ‘The CX3CL1-CX3CR1 chemokine axis can contribute to tumor immune evasion and blockade with a novel CX3CR1 monoclonal antibody enhances response to anti-PD-1 immunotherapy’, Front. Immunol., vol. 14, Sep. 2023, doi: 10.3389/fimmu.2023.1237715.

[26] Chen, X. Zhou, Y. Fan, and C. Wang, ‘Identification and validation of prognostic biomarkers related to tumor immune invasion in pancreatic cancer’, Front. Genet., vol. 16, Mar. 2025, doi: 10.3389/fgene.2025.1556544.

[27] G. Nardone et al., ‘YAP regulates cell mechanics by controlling focal adhesion assembly’, Nat. Commun., vol. 8, no. 1, p. 15321, May 2017, doi: 10.1038/ncomms15321.

[28] Liu, Y. Song, D. Li, and B. Wang, ‘Regulation of the tumor immune microenvironment by the Hippo Pathway: Implications for cancer immunotherapy’, Int. Immunopharmacol., vol. 122, p. 110586, Sep. 2023, doi: 10.1016/j.intimp.2023.110586.

[29] H. Buckarma et al., ‘The YAP-Interacting Phosphatase SHP2 Can Regulate Transcriptional Coactivity and Modulate Sensitivity to Chemotherapy in Cholangiocarcinoma’, Mol. Cancer Res., vol. 18, no. 10, pp. 1574–1588, Oct. 2020, doi: 10.1158/1541-7786.MCR-20-0165.

[30] Baroja, N. C. Kyriakidis, G. Halder, and I. M. Moya, ‘Expected and unexpected effects after systemic inhibition of Hippo transcriptional output in cancer’, Nat. Commun., vol. 15, no. 1, p. 2700, Mar. 2024, doi: 10.1038/s41467-024-46531-1.

[31] Y. Xiao and J. Dong, ‘The Hippo Signaling Pathway in Cancer: A Cell Cycle Perspective’, Cancers, vol. 13, no. 24, p. 6214, Dec. 2021, doi: 10.3390/cancers13246214.

[32] K. Ostacolo et al., ‘ATG7(2) Interacts With Metabolic Proteins and Regulates Central Energy Metabolism’, Traffic, vol. 25, no. 4, p. e12933, 2024, doi: 10.1111/tra.12933.

[33] A. Lee et al., ‘Autophagy is a gatekeeper of hepatic differentiation and carcinogenesis by controlling the degradation of Yap’, Nat. Commun., vol. 9, no. 1, p. 4962, Nov. 2018, doi: 10.1038/s41467-018-07338-z.

[34] Y. Nishida et al., ‘Discovery of Atg5/Atg7-independent alternative macroautophagy’, Nature, vol. 461, no. 7264, pp. 654–658, Oct. 2009, doi: 10.1038/nature08455.

[35] O. Aguilera, L. R. Delgui, F. Reggiori, P. S. Romano, and M. I. Colombo, ‘Autophagy as an innate immunity response against pathogens: a Tango dance’, FEBS Lett., vol. 598, no. 1, pp. 140–166, 2024, doi: 10.1002/1873-3468.14788.

[36] S. Upadhyay and J. A. Philips, ‘LC3-associated phagocytosis: host defense and microbial response’, Curr. Opin. Immunol., vol. 60, pp. 81–90, Oct. 2019, doi: 10.1016/j.coi.2019.04.012.

[37] L. Heckmann, E. Boada-Romero, L. D. Cunha, J. Magne, and D. R. Green, ‘LC3-Associated Phagocytosis and Inflammation’, J. Mol. Biol., vol. 429, no. 23, pp. 3561–3576, Nov. 2017, doi: 10.1016/j.jmb.2017.08.012.

[38] W. Hou et al., ‘Inhibiting autophagy selectively prunes dysfunctional tumor vessels and optimizes the tumor immune microenvironment’, Theranostics, vol. 15, no. 1, pp. 258–276, Jan. 2025, doi: 10.7150/thno.98285.

[39] Chen, Y. Liang, S. Hu, J. Jiang, M. Zeng, and M. Luo, ‘Role of ATG7-dependent non-autophagic pathway in angiogenesis’, Front. Pharmacol., vol. 14, Jan. 2024, doi: 10.3389/fphar.2023.1266311.

[40] S. Xu et al., ‘Targeting immune checkpoints on tumor-associated macrophages in tumor immunotherapy’, Front. Immunol., vol. 14, p. 1199631, 2023, doi: 10.3389/fimmu.2023.1199631.

[41] A. Alturki, ‘Review of the Immune Checkpoint Inhibitors in the Context of Cancer Treatment’, J. Clin. Med., vol. 12, no. 13, p. 4301, Jun. 2023, doi: 10.3390/jcm12134301.

[42] P. Sharma et al., ‘Immune checkpoint therapy-current perspectives and future directions’, Cell, vol. 186, no. 8, pp. 1652–1669, Apr. 2023, doi: 10.1016/j.cell.2023.03.006.

[43] E. Karathanasis and K. B. Ghaghada, ‘Crossing the barrier: Treatment of brain tumors using nanochain particles’, Wiley Interdiscip. Rev. Nanomed. Nanobiotechnol., vol. 8, no. 5, pp. 678–695, Sep. 2016, doi: 10.1002/wnan.1387.

[44] Zhou, X. Chen, J. Cao, and H. Gao, ‘Overcoming the biological barriers in the tumor microenvironment for improving drug delivery and efficacy’, J. Mater. Chem. B, vol. 8, no. 31, pp. 6765–6781, Aug. 2020, doi: 10.1039/D0TB00649A.

[45] Sakurai, T. Hada, S. Yamamoto, A. Kato, W. Mizumura, and H. Harashima, ‘Remodeling of the Extracellular Matrix by Endothelial Cell-Targeting siRNA Improves the EPR-Based Delivery of 100 nm Particles’, Mol. Ther., vol. 24, no. 12, pp. 2090–2099, Dec. 2016, doi: 10.1038/mt.2016.178.

[46] Zhu, F. Perche, T. Wang, and V. P. Torchilin, ‘Matrix metalloproteinase 2-sensitive multifunctional polymeric micelles for tumor-specific co-delivery of siRNA and hydrophobic drugs’, Biomaterials, vol. 35, no. 13, pp. 4213–4222, Apr. 2014, doi: 10.1016/j.biomaterials.2014.01.060.

[47] L. Mainz et al., ‘Autophagy Blockage Reduces the Incidence of Pancreatic Ductal Adenocarcinoma in the Context of Mutant Trp53’, Front. Cell Dev. Biol., vol. 10, p. 785252, 2022, doi: 10.3389/fcell.2022.785252.

[48] H. Lee et al., ‘Atg7 modulates p53 activity to regulate cell cycle and survival during metabolic stress’, Science, vol. 336, no. 6078, pp. 225–228, Apr. 2012, doi: 10.1126/science.1218395.

