## Supplemental figures for "ATG7 function promotes pancreatic cancer progression independently of autophagy"

A

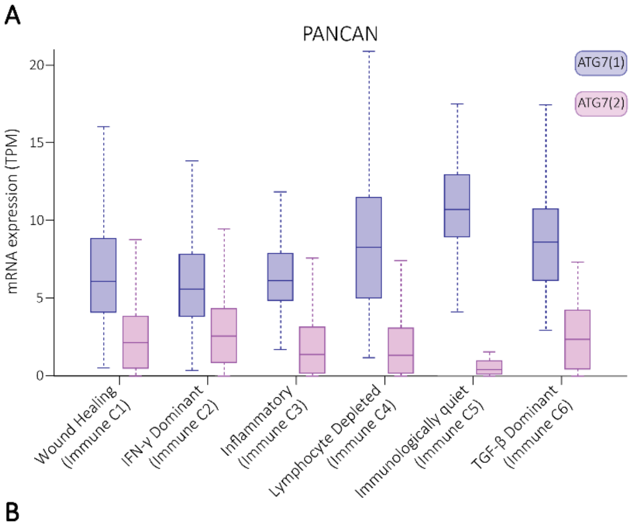


| | ATG7(1) | ATG7(2) |
|---|---|---|
| C1 vs. C2 | * | * |
| C1 vs. C3 | ns | * |
| C1 vs. C4 | **** | ** |
| C1 vs. C5 | **** | **** |
| C1 vs. C6 | **** | ns |
| C2 vs. C3 | ns | **** |
| C2 vs. C4 | **** | **** |
| C2 vs. C5 | **** | **** |
| C2 vs. C6 | **** | ns |
| C3 vs. C4 | **** | ns |
| C3 vs. C5 | **** | * |
| C3 vs. C6 | **** | ns |
| C4 vs. C5 | *** | * |
| C4 vs. C6 | ns | ns |
| C5 vs. C6 | ns | ** |

B

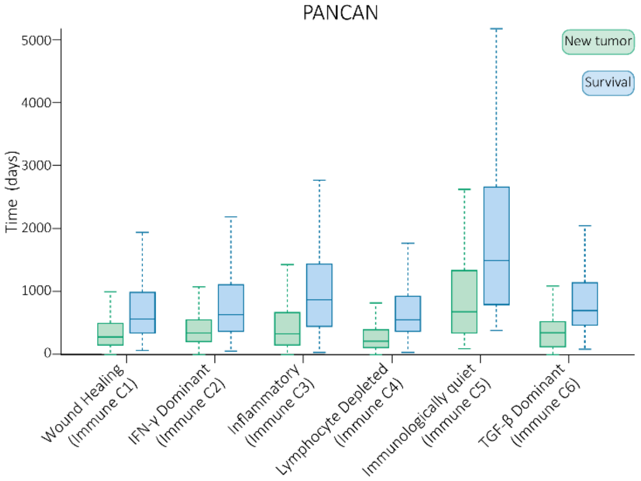


| | Survival | New Tumour |
|---|---|---|
| C1 vs. C2 | ns | ns |
| C1 vs. C3 | ** | ns |
| C1 vs. C4 | ns | ** |
| C1 vs. C5 | **** | **** |
| C1 vs. C6 | ns | ns |
| C2 vs. C3 | * | ns |
| C2 vs. C4 | * | **** |
| C2 vs. C5 | **** | **** |
| C2 vs. C6 | ns | ns |
| C3 vs. C4 | *** | **** |
| C3 vs. C5 | **** | **** |
| C3 vs. C6 | ns | ns |
| C4 vs. C5 | **** | **** |
| C4 vs. C6 | ns | ns |
| C5 vs. C6 | **** | **** |

C

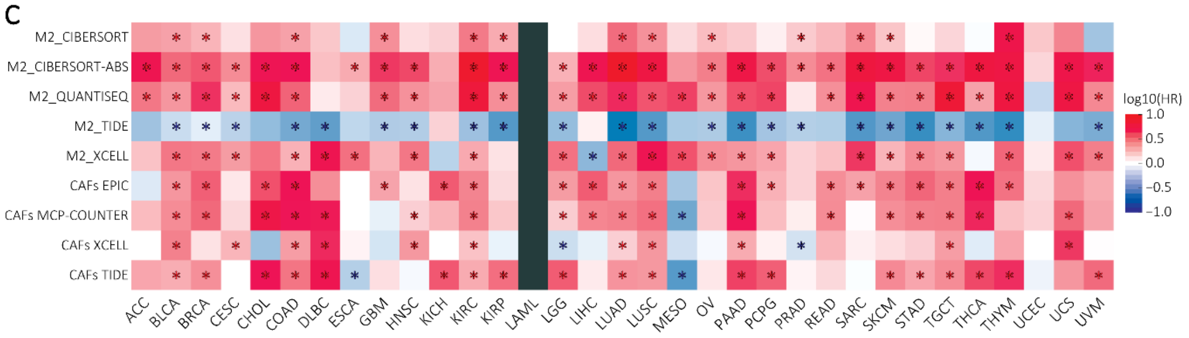


**Figure S1: ATG7(2) is linked with immune modulation in cancer.**

**A:** Expression plot of ATG7(1) and ATG7(2) in different immune subtypes of cancer samples. **B:** Box plot of survival time or time before new tumour event in the same samples. **A-B:** One way ANOVA was performed between the different groups, with *: $p < 0.05$; **: $p < 0.01$; ***: $p < 0.001$; ****: $p < 0.0001$. **C:** Correlation of ATG7 expression with immune cells infiltration across cancers. Red: positive correlation between ATG7 expression and tumour infiltration; Blue: negative correlation between ATG7 expression and tumour infiltration; *: significant correlation $p < 0.05$.

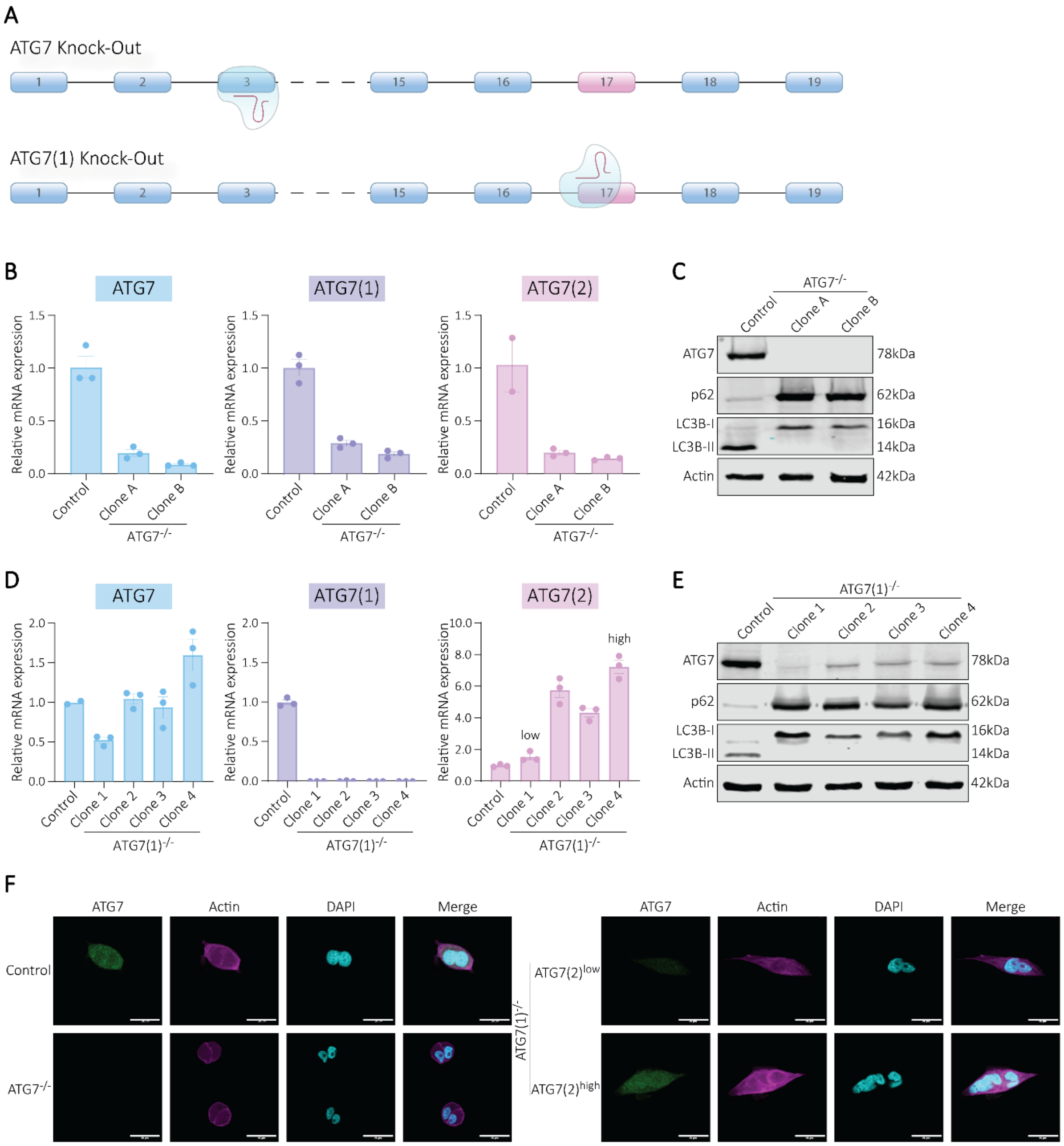


**Figure S2: ATG7(2) is localized in the cell cytoplasm and fails to initiate autophagy.**
**A:** Schematic of the CRISPR/Cas9 gRNAs targeting exon 3 for ATG7 knock-out; or exon 17 for ATG7(1) knock-out. **B:** mRNA quantification of ATG7, ATG7(1) and ATG7(2) using qPCR in Capan-1 control and ATG7$^{-/-}$ cells. **C:** Western-Blot analysis of Capan-1 control and ATG7$^{-/-}$ cells. **D:** mRNA quantification of ATG7, ATG7(1) and ATG7(2) using qPCR in Capan-1 control and ATG7(1)$^{-/-}$ cells. **E:** Western-Blot analysis of Capan-1 control and ATG7(1)$^{-/-}$ cells. **F:** Immunostaining of Capan-1 control, ATG7$^{-/-}$ and ATG7(1)$^{-/-}$ cells, imaged under a confocal microscope, scale bar: 25µm.

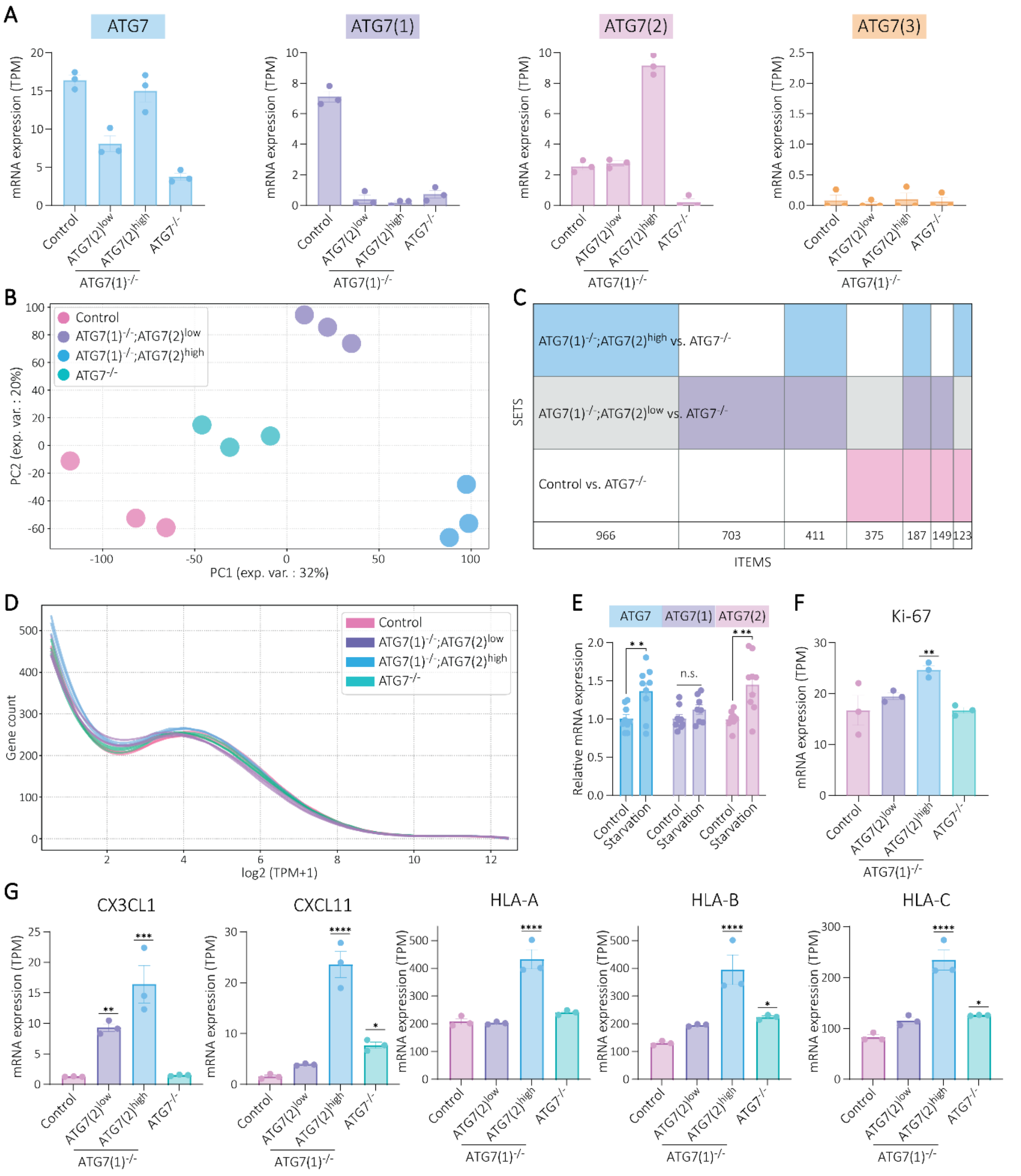


**Figure S3: ATG7(2) regulates the expression of numerous genes in PAAD cells.**
**A:** mRNA expression of *ATG7*, *ATG7(1)*, *ATG7(2)* and *ATG7(3)* in control, ATG7(1)$^{-/-}$ and ATG7$^{-/-}$ cells. **B:** PCA plot of gene expression in control, ATG7(1)$^{-/-}$ and ATG7$^{-/-}$ cells. **C:** Supervenn diagram of control vs ATG7$^{-/-}$; ATG7(1)$^{-/-}$;ATG7(2)$^{low}$ vs ATG7$^{-/-}$ and ATG7(1)$^{-/-}$;ATG7(2)$^{high}$ vs ATG7$^{-/-}$ differentially expressed genes. **D:** Distribution of transcriptome profiles. Each line represents a sample. **E:** mRNA quantification of *ATG7*, *ATG7(1)* and *ATG7(2)* using qPCR in Capan-1 control cells after 4h in complete growth medium or glucose-depleted growth medium. **F:** mRNA expression of the proliferation marker *Ki-67* in Capan-1 control, ATG7(1)$^{-/-}$ and ATG7$^{-/-}$ cells, from RNA sequencing. **G:** mRNA expression of cytokines and components of the MHC-I system, in Capan-1 control, ATG7(1)$^{-/-}$ and ATG7$^{-/-}$ cells, from RNA sequencing. **E:** Multiple t-tests were performed between the control and the starvation condition / **F-G:** One way ANOVA was performed between the control and every other condition, with *: $p < 0.05$; **: $p < 0.01$; ***: $p < 0.001$; ****: $p < 0.0001$.

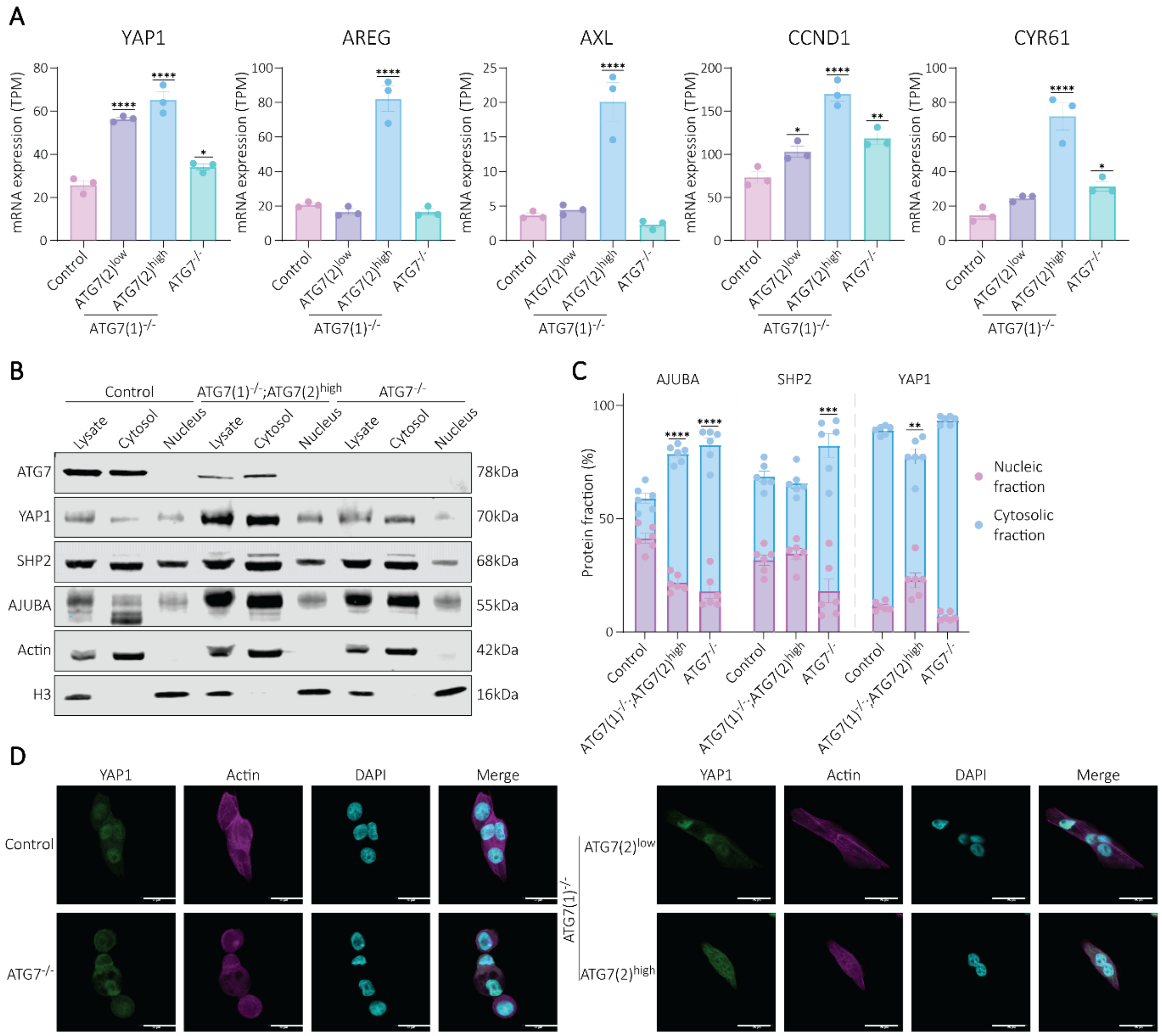


**Figure S4: ATG7(2) regulates YAP activity in PAAD cells.**
**A:** mRNA expression of YAP target genes in control, ATG7(1)$^{-/-}$ and ATG7$^{-/-}$ cells, from RNA sequencing. **B:** Western-blotting of YAP1, SHP2 and AJUBA in nuclear and cytoplasmic fractions of Capan-1 control, ATG7(1)$^{-/-}$ and ATG7$^{-/-}$ cells. **C:** Quantification of the cytosolic and nuclear fractions, in %, for AJUBA, SHP2 and YAP1. **A,C:** One way ANOVA was performed between the control and every other condition, with *: $p < 0.05$; **: $p < 0.01$; ***: $p < 0.001$; ****: $p < 0.0001$. **D:** Immunostaining of Capan-1 control, ATG7$^{-/-}$ and ATG7(1)$^{-/-}$ cells stained for YAP cellular localization, imaged under a confocal microscope, scale bar: 25µm.

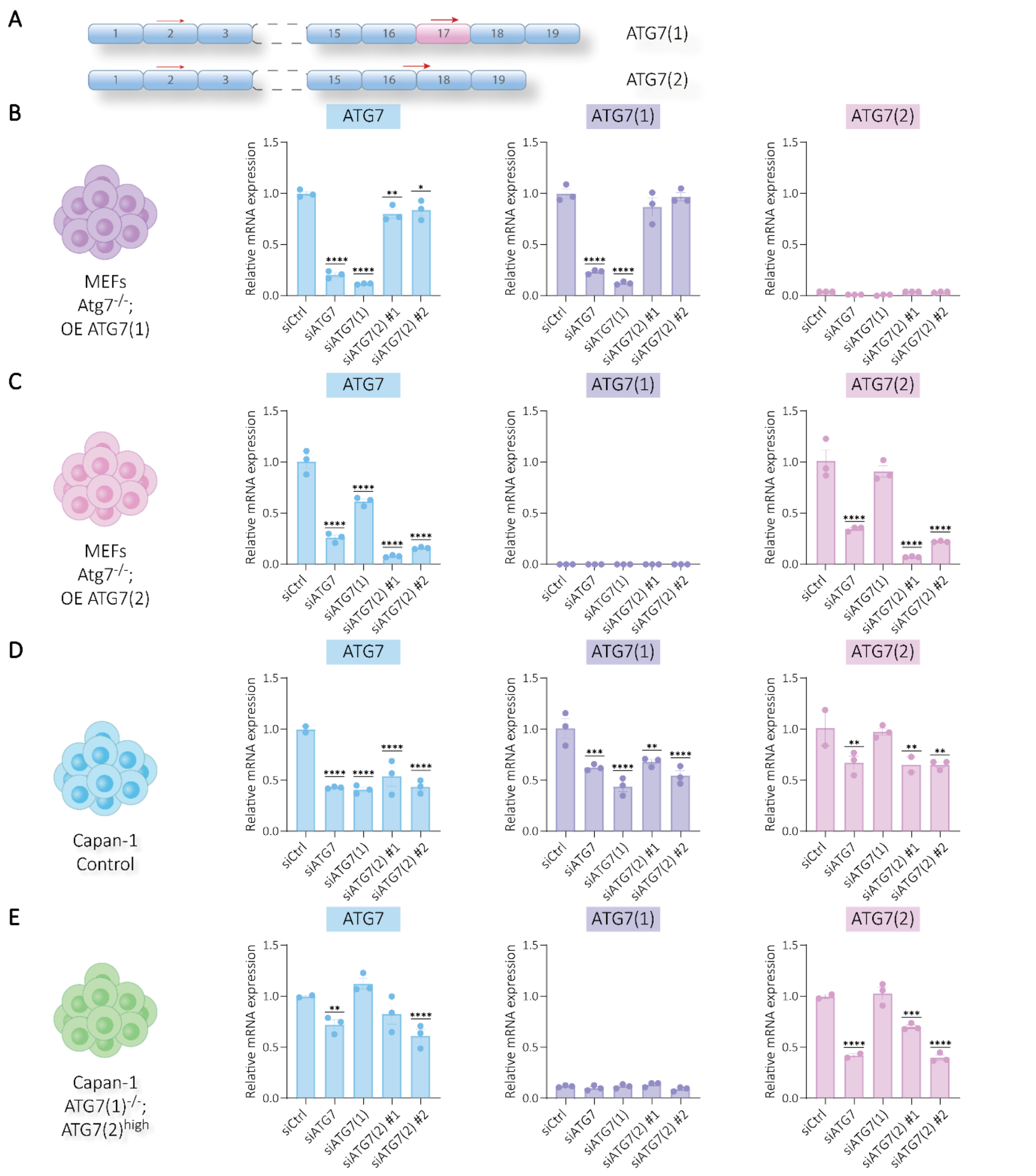


**Figure S5: siRNA efficiently knocks down ATG7(2) in a specific manner.**
**A:** Schematic of the siRNA sequences targeting ATG7(2) specifically. **B:** mRNA expression of *ATG7* isoforms using qPCR in MEF Atg7$^{-/-}$ cells overexpressing ATG7(1) after siATG7(2) or control siRNA treatment for 48h. **C:** mRNA expression of *ATG7* isoforms using qPCR in MEF Atg7$^{-/-}$ cells overexpressing ATG7(2) after siATG7(2) or control siRNA treatment for 48h. **D:** mRNA expression of *ATG7* isoforms using qPCR in Capan-1 control cells after siATG7(2) or control siRNA treatment for 72h. **E:** mRNA expression of *ATG7* isoforms using qPCR in Capan-1 ATG7(1)$^{-/-}$;ATG7(2)$^{high}$ cells after siATG7(2) or control siRNA treatment for 72h. **B-E:** One way ANOVA was performed between the control and the every other condition, with *: $p < 0.05$; **: $p < 0.01$; ***: $p < 0.001$; ****: $p < 0.0001$.

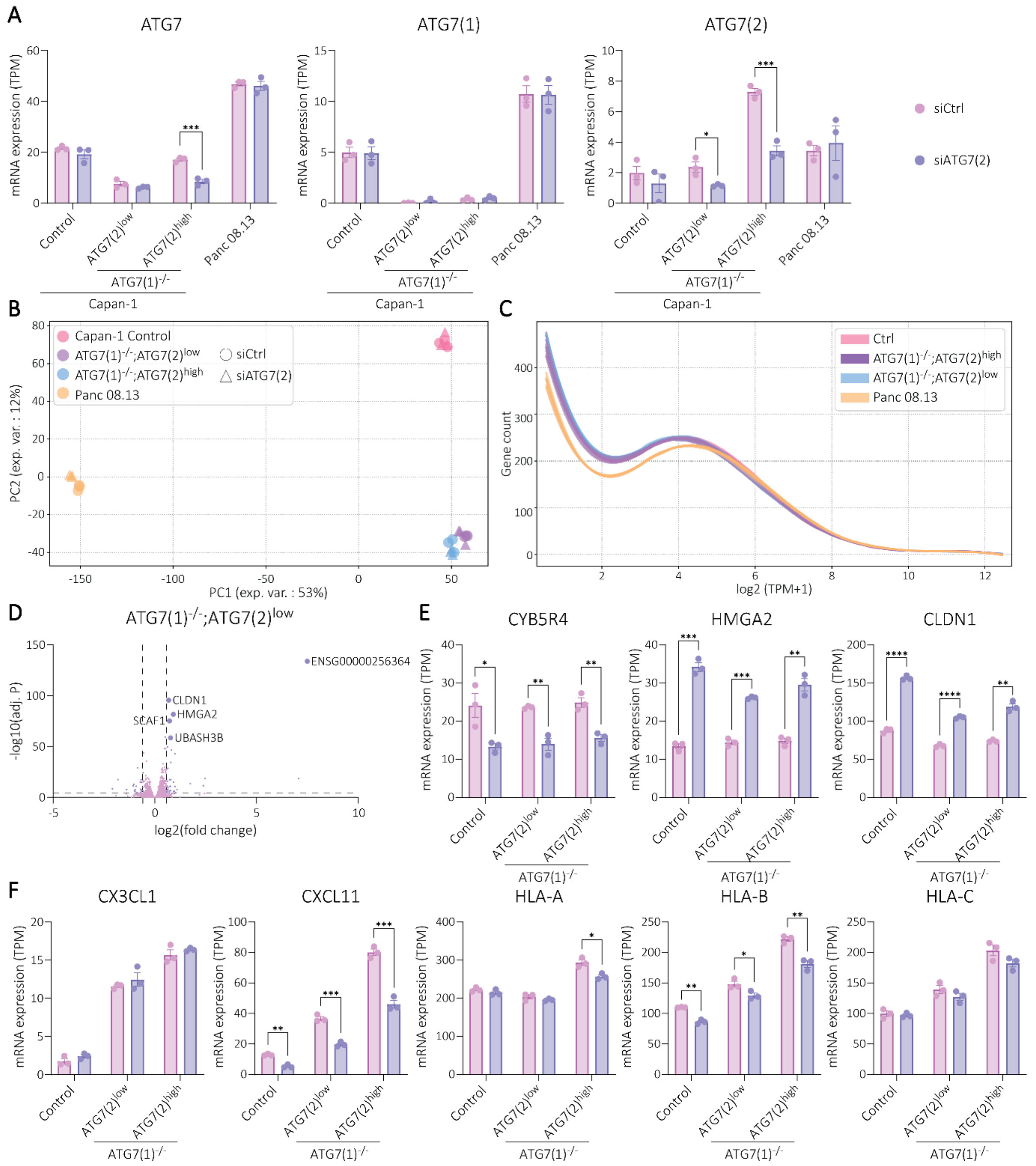


**Figure S6: ATG7(2) inhibition affects gene expression in PAAD cells.**

**A:** mRNA expression of *ATG7*, *ATG7(1)*, *ATG7(2)* and *ATG7(3)* in in Panc 08.13 and Capan-1 Control, ATG7(1)$^{-/-}$;ATG7(2)$^{low}$ and ATG7(1)$^{-/-}$;ATG7(2)$^{high}$ cells, treated with control siRNA or siATG7(2), n=3. **B:** PCA plot of gene expression in Panc 08.13 and Capan-1 Control, ATG7(1)$^{-/-}$;ATG7(2)$^{low}$ and ATG7(1)$^{-/-}$;ATG7(2)$^{high}$ cells, treated with control siRNA or siATG7(2), n=3. **C:** Distribution of transcriptome profiles. Each line represents a sample. **D:** Volcano plots depicting differentially expressed genes in Capan-1 ATG7(1)$^{-/-}$;ATG7(2)$^{low}$ cells when treated with siATG7(2) for 72h. Pink dots represent genes that are not differentially expressed, while purple dots represent genes that are differentially expressed. The 5 genes with the highest difference in expression are labelled. **E:** mRNA expression of *CYB5R4*, *HMGA2* and *CLDN1* in Capan-1 Control, ATG7(1)$^{-/-}$;ATG7(2)$^{low}$ and ATG7(1)$^{-/-}$;ATG7(2)$^{high}$ cells, treated with control siRNA or siATG7(2), n=3. **F:** mRNA expression of cytokines and components of the MHC-I system, in Capan-1 Control, ATG7(1)$^{-/-}$;ATG7(2)$^{low}$ and ATG7(1)$^{-/-}$;ATG7(2)$^{high}$ cells, treated with control siRNA or siATG7(2), n=3. **A,E-F:** Multiple t-tests were performed between the control siRNA and the siATG7(2) condition, with *: $p < 0.05$; **: $p < 0.01$; ***: $p < 0.001$; ****: $p < 0.0001$.

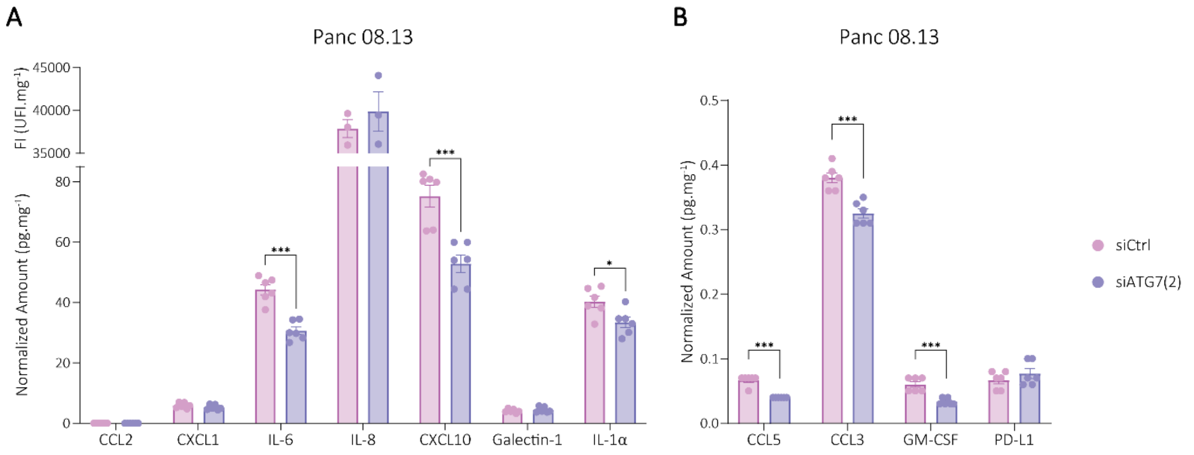


**Figure S7: ATG7(2) inhibition affects immune signalling in PAAD cells.**

**A-B:** Protein secretion of factors involved in in T cells, Neutrophils and MDSCs recruitment and polarization, after siATG7(2) or control siRNA treatment for 72h in Panc 08.13 cells. From Multiplex Luminex® quantification. Multiple t-tests were performed between the control and the siATG7(2) or siYAP1 condition, with *: $p < 0.05$; **: $p < 0.01$; ***: $p < 0.001$; ****: $p < 0.0001$.

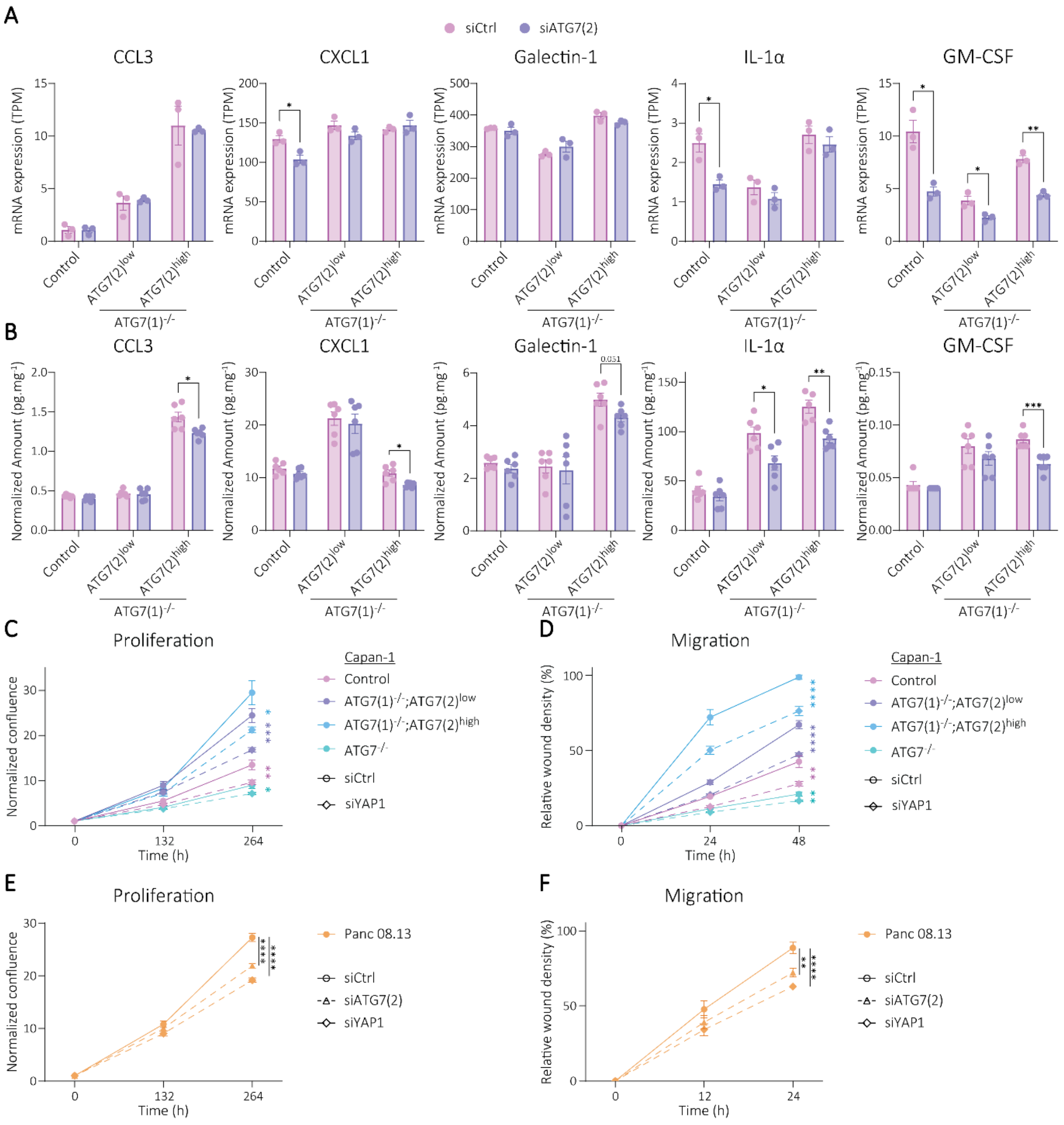


**Figure S8: YAP1 inhibition reduces PAAD cell migration and proliferation *in vitro*, and ATG7(2) inhibition affects immune signalling.**

**A:** mRNA expression of factors involved in in T cells, Neutrophils and MDSCs recruitment and polarization, after siATG7(2) or control siRNA treatment for 72h. From RNA sequencing. **B:** Protein secretion of factors involved in in T cells, Neutrophils and MDSCs recruitment and polarization, after siATG7(2) or control siRNA treatment for 72h. From Multiplex Luminex® quantification. **C:** Normalized proliferation ratios of Capan-1 control and ATG7(1)$^{-/-}$ cells after siYAP1 or control siRNA treatment every 72h, at 132h and 264h timepoints after seeding. **D:** Normalized migration ratios of Capan-1 control and ATG7(1)$^{-/-}$ cells after siYAP1 or control siRNA treatment every 72h, at 24h and 48h timepoints after wound making. **E:** Normalized proliferation ratios of Panc 08.13 cells after siYAP1, siATG7(2) or control siRNA treatment every 72h, at 132h and 264h timepoints after seeding. **F:** Normalized migration ratios of Panc 08.13 cells after siYAP1, siATG7(2) or control siRNA treatment every 72h, at 24h and 48h timepoints after wound making. **A-F:** Multiple t-tests were performed between the control and the siATG7(2) or siYAP1 condition, with *: $p < 0.05$; **: $p < 0.01$; ***: $p < 0.001$; ****: $p < 0.0001$.

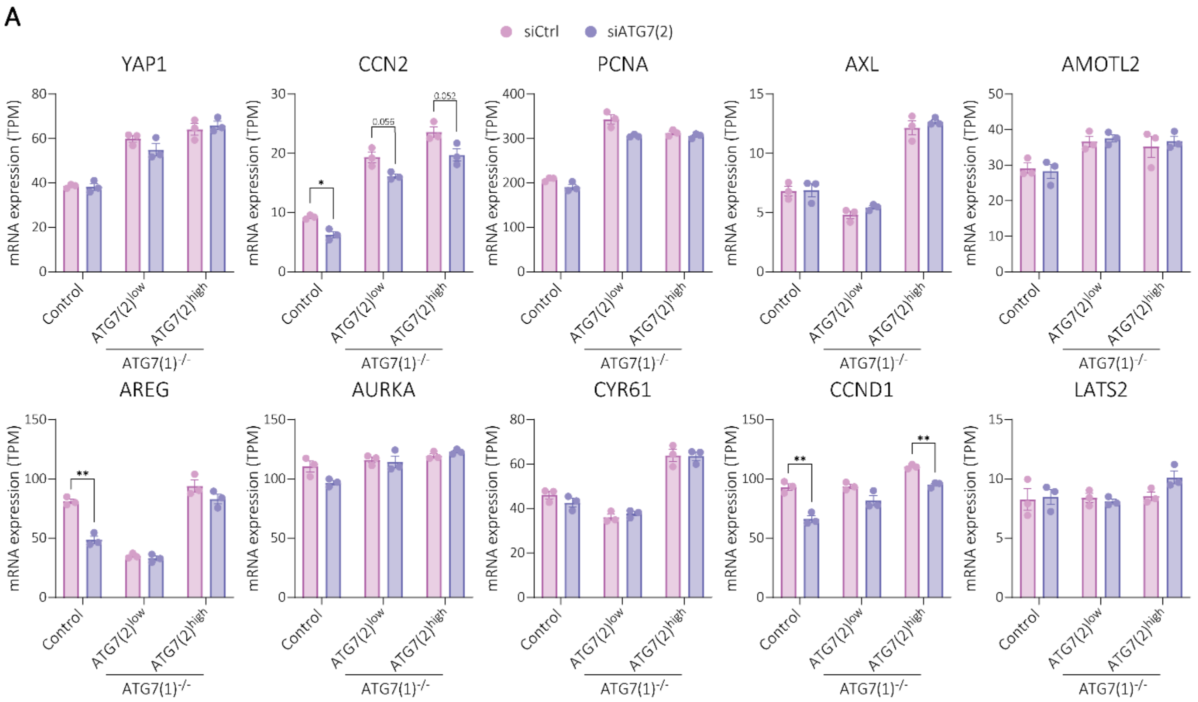


**Figure S9: ATG7(2) inhibition does not significantly impact YAP1 activity.**
**A:** mRNA expression of YAP target genes after siATG7(2) or control siRNA treatment for 72h in Capan-1 Control, ATG7(1)$^{-/-}$ and ATG7$^{-/-}$ cells. From RNA sequencing. Multiple t-tests were performed between the control and every other condition, with *: $p < 0.05$; **: $p < 0.01$; ***: $p < 0.001$; ****: $p < 0.0001$.

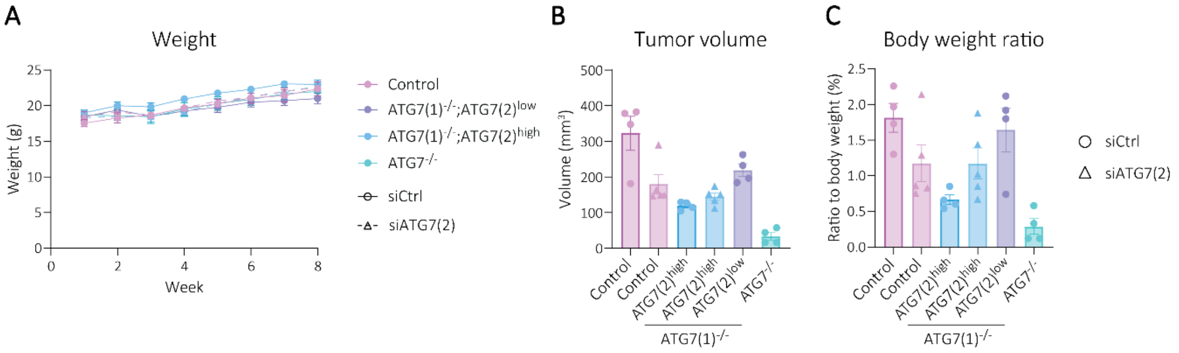


**Figure S10: ATG7(2) contributes to PAAD progression *in vivo*.**
**A:** Evolution of the body weight in Capan-1 Control, ATG7(1)$^{-/-}$ and ATG7$^{-/-}$ cohorts treated with control siRNA or siATG7(2) (n=5). **B:** Tumour volumes at the end of the study. **C:** Ratios of tumour weight to body weight at the end of the study.
